# Genome and Genetic Engineering of the House Cricket (*Acheta domesticus*): Applications for Sustainable Agriculture

**DOI:** 10.1101/2022.12.14.520443

**Authors:** Aaron T. Dossey, Brenda Oppert, Fu-Chyun Chu, Marcé D. Lorenzen, Brian Scheffler, Sheron Simpson, Sergey Koren, J. Spencer Johnston, Kosuke Kataoka, Keigo Ide

**Affiliations:** All Things Bugs LLC; 2211 Snapper Ln., Oklahoma City, OK 73130; USDA Agricultural Research Service, Center for Grain and Animal Health Research; 1515 College, Ave., Manhattan, KS 66502 USA; Department of Entomology and Plant Pathology, North Carolina State University, Raleigh, NC 27695 USA; USDA Agricultural Research Service, Jamie Whitten Delta States Research Center; 141 Experiment Station Road, Stoneville, MS 38776-0036 USA; Genome Informatics Section, Computational and Statistical Genomics Branch, National Human Genome Research Institute, National Institutes of Health, Bethesda, MD, 20894 USA; Department of Entomology, Texas A&M University, College Station, TX 77843, USA; Faculty of Science and Engineering, Waseda University; 2-2 TWIns #02C214, Wakamatsu-cho, Shinjuku-ku, Tokyo, 162-8480, JAPAN

**Keywords:** Food Security, Cricket, Protein, Genome, Sustainability, Agriculture, Insects as food and feed, *Acheta domesticus*, Insect Genome, CRISPR, CRPSIR Cas9, Genetic engineering

## Abstract

The house cricket, *Acheta domesticus*, is one of the most farmed insects worldwide and the foundation of an emerging industry for the use of insects as a sustainable food source. Edible insects present a promising alternative for protein production amid a plethora of recent reports on climate change and biodiversity loss largely driven by agriculture. As with other agricultural crops, genetic resources are needed to improve crickets for food and other applications. We present the first high quality annotated genome assembly of *A. domesticus* which was assembled from long read data and scaffolded to chromosome level from long range data, providing information on promoters and genes needed for genetic manipulation. Gene groups that may be useful for improving the value of these insects to farmers were manually annotated, mainly genes related to immunity. Metagenome scaffolds in the *A. domesticus* assembly, including those from bacteria, other microbes and viruses such as Invertebrate Iridescent Virus 6 (IIV6), were submitted in a separate accession as host-associated sequences. We demonstrate both CRISPR/Cas9-mediated knock-in and knock-out of selected genes and discuss implications for the food, pharmaceutical and other industries. RNAi was demonstrated to disrupt the function of the *vermilion* eye-color gene to produce a useful white-eye biomarker phenotype. We are utilizing these data to develop base technologies and methodologies for downstream commercial applications, including the generation of more nutritious and disease resistant crickets as well as lines producing valuable bioproducts such as vaccines and antibiotics. We also discuss how this foundational research can play a critical role in utilizing the largest, most diverse yet almost entirely untapped biological resource on Earth: Class Insecta.

**Significance Statement:** Sequencing and assembly of the genome of the house cricket has led to improvements in farmed insects for food, pharmaceutical and other applications.

## Introduction

Multiple reports demonstrate that destruction of natural habitats and pollution from human activity, largely a result of land clearing for agriculture and climate change, have led to substantial global biodiversity loss and mass extinction [1–4]. Threats to biodiversity are also threats to all life on earth, including to humans [4–7]. With historic levels of biodiversity loss and an increasing human population, it is critical to reduce consumption of natural resources from earth and its ecosphere. The UN expects the human population to grow to nearly 10 billion by 2050 [8], and food demand is projected to increase up to 62% [9]. Climate change, reduced productivity of agricultural lands, overfishing, dwindling freshwater, pollution from fertilizers and pesticides, and a host of other factors resulting from population increase will place a disproportionate burden on Earth’s ecosphere. Population increases and scarcity of natural resources and food will lead to conflict around the world [10]. Already nearly half of land on earth is used for agriculture and at least 70% of agricultural land, 30% of the land on earth, is used for livestock [10–13]. Meanwhile, global market demand for protein continues to grow rapidly from $38 billion in 2019 [14] and to a projected increase of $ 32.56 billion by 2028 [15]. Unfortunately, expanding the amount of land used for livestock production is neither feasible nor sustainable.

The value of insects to life on earth, including humans, cannot be over-stated [16]. Insects provide critical ecological services including pollination and nutrient cycling which support practically all life on earth [17–21]. Not only do insects provide important functions for the natural world, they also are a nearly entirely untapped bioresource for humans. Insects contain a wealth of chemical biodiversity [18, 22–23]. For example, insects are a promising source of high-quality protein as well as oils, chitin nutrients and other bioproducts with a substantially lower environmental footprint than other livestock [24–25]. Utilization of insects in food in place of less sustainable protein sources, such as vertebrate livestock, will significantly reduce the human impact on the environment, including biodiversity loss, climate change and clean water depletion.

It has been well established that insects are a dense source of dietary nutrients [26–27]. Recent research demonstrates evidence of other health benefits of consuming insects including biomarkers for improved gut health [28]. Zoonotic diseases, such as viruses, are much less likely to jump from farm-raised insects to humans than from mammals or bird livestock due to large genetic differences between humans and arthropods [25, 29–30]. Additionally, studies show that foodborne pathogen loads, such as *Salmonella spp*. and *Listeria monocytogenes*, in farmed insects are low and often absent [31–32]. The European Commission recently approved the focus species of this paper, *A. domesticus* as well as *Tenebrio molitor* (yellow mealworm) as a safe novel food [33].

Given the excellent nutritional value of insects and their ability to be grown efficiently in small confined spaces with minimal resource inputs, they could be a critically important solution for sustainable food/protein production [24-25, 34-36]. Entomophagy, consumption of insects as food, is accepted and practiced by over 2 billion people worldwide [37–39]. Some attractive features offered by insects include low land, water, and natural resource utilization [34], reduced greenhouse gas emissions [36, 40], highly prolific (1,500-3,000 eggs per female) [34, 41–42] and short life cycles for rapid scale-up to support food security. Insects are amenable to modularized vertical farming as well as automated and urban farming. They also can be grown closer to cities and/or processing facilities. Due to their small size and growing efficiencies, insects are likely the only animal feasible for production in space exploration such as at stations on the moon and Mars [43–44].

In addition to improved production efficiencies with lower environmental impact, diversification of our food supply is critical for food security. According to the FAO, 75% of food comes from only 12 plant and 5 animal species [45–46]. Class Insecta is the largest, most diverse group of organisms on Earth, with approximately 950,000 species described and 4-30 million estimated [18, 47–48]. At least 2,000 have been identified as eaten around the world and many can be farmed [37-38, 24-25, 49]. One of the food security benefits of biodiversity offered by insects is the ability to rapidly switch species in response to crop loss from disease [25, 50].

In the past decade an insect-based food and feed industry has emerged [51–52]. This industry is primarily based on mass production of crickets (particularly *A. domesticus* and *Grylloides signalus*) for human food [25, 53]. As of 2012, at least 2,000 tons of crickets are produced annually in the US (over 2 billion adult crickets) [50–51]. While eating insects has been promoted in the past [29, 35, 38, 49, 53–56], this new industry offers long term-term solutions. By 2027, the edible insect market is projected to reach $4.63 billion, increasing 26.5% per year [15]. Major investments of as much as $400 million for a single insect farming company have been made [57–59]. These are some of the largest agricultural investments in the world.

Insect genomes are critical to all fields of biological science, from medicine to agriculture and conservation [60], with the *Drosophila melanogaster* genome as one of the first model organisms with a reference genome assembly [61]. Mass produced insects can contribute positively to sustainable human existence but require the development of genetic resources including reference quality genome assemblies. [62]. Not only is sequencing more affordable, improvements in long read length and accuracy as well as scaffolding technology have resulted in near complete genome assemblies, often assembled to chromosomal level [63–65]. Food and agricultural systems have benefitted from sequencing, including vaccine manufacturing, genetic engineering of crops playing a critical role [66]. The short lifespan and high fecundity of insects offers an opportunity to use genetic modification with more rapid success rates than with larger animals and many plant crops to produce not only food but bioproduction of other valuable materials such as vaccine antigens. Coupling genome research and CRISPR/Cas9-mediated genome editing offers an unprecedented potential to rapidly improve insect crops such as increasing nutritional content, reducing allergenicity, providing resistance to diseases, biomanufacturing, improving feed conversion, increasing growth rate, reducing mortality and other attributes desired by farmers [67–68]. More studies on insect biology in the context of genome sequences are critically important to not only investigate these species while we still can, but also to inspire the world to make greater strides in conserving and preserving biodiversity and natural habitats by demonstrating their value [16].

To provide genetic resources to the insect as food industry, we present the first high quality annotated genome assembly of *A. domesticus* and demonstrate how genetics can be used to address limitations in these insects as a food resource. The assembly is scaffolded to chromosome level, providing information on promoters and genes needed for genetic transformation. We demonstrate both CRISPR/Cas9-mediated knock-in and knock-out of genes, and discuss implications for the food, biomedical, vaccine and other bioproduction industries.

Specific data includes: 1) metrics of a high quality chromosomal level long read and scaffolded genome assembly, 2) results utilizing CRISPR/Cas9 technology to genetically modify (knock-in and knock-out) genes in this cricket species (to our knowledge the first published genetically engineered lines of this species), 3) demonstration of successful use of RNAi to reduce gene expression and 4) annotation of a select group of genes in our genomic data (including genes from cricket related to immunity as well as scaffolded microbe and virus sequences).

Subsequently, we provide a discussion on how this work utilizing modern cutting edge genomic research can play a critical role in utilizing the largest, most diverse yet almost entirely untapped biological resource on Earth: Class Insecta.

## Results

### Genome Size and Assembly

The size of the *A. domesticus* genome was measured by flow cytometry as described in [69]. The genome of female *A. domesticus* was larger than males (2,378.8 ± 9.7 vs. 2,149.6 ± 11.2 Mb, respectively, Table S1). The size of the X-chromosome was estimated as 387 Mb, the difference between the 2C genome size measurement of the XX female and the XO male. The average size of autosomal chromosomes was 177 Mb.

Genomic DNA was extracted from a single *A. domesticus* adult male for long read sequencing, and nuclear DNA from the same adult male was used for long range data in scaffolding. The draft assembly of long read data contained 28,340 contigs with an N_50_ of 321,088 (Table 1). The final assembly with long range (Chicago and Hi-C) data contained 16,290 scaffolds originally with an N_50_ of 221.839 Mb and a BUSCO score of 94.8%. After contaminant scaffolds (metagenome, mitochondrial, duplicate, and vector) were removed, the final *A. domesticus* genome assembly contained 9,964 scaffolds consisting of 2,346,604,983 bp, comparable to the predicted size of 2.150 Gb. The final assembly had 11 larger chromosomes that totaled 2,137,583,504 bases, presumably representing the 10 autosomal chromosomes and single X sex chromosome predicted for males (Figure 1). The longest scaffold in the assembly was 304 Mb (scaffold_14394) and may represent a portion of the X chromosome (predicted as 387 Mb).

**Figure 1.**
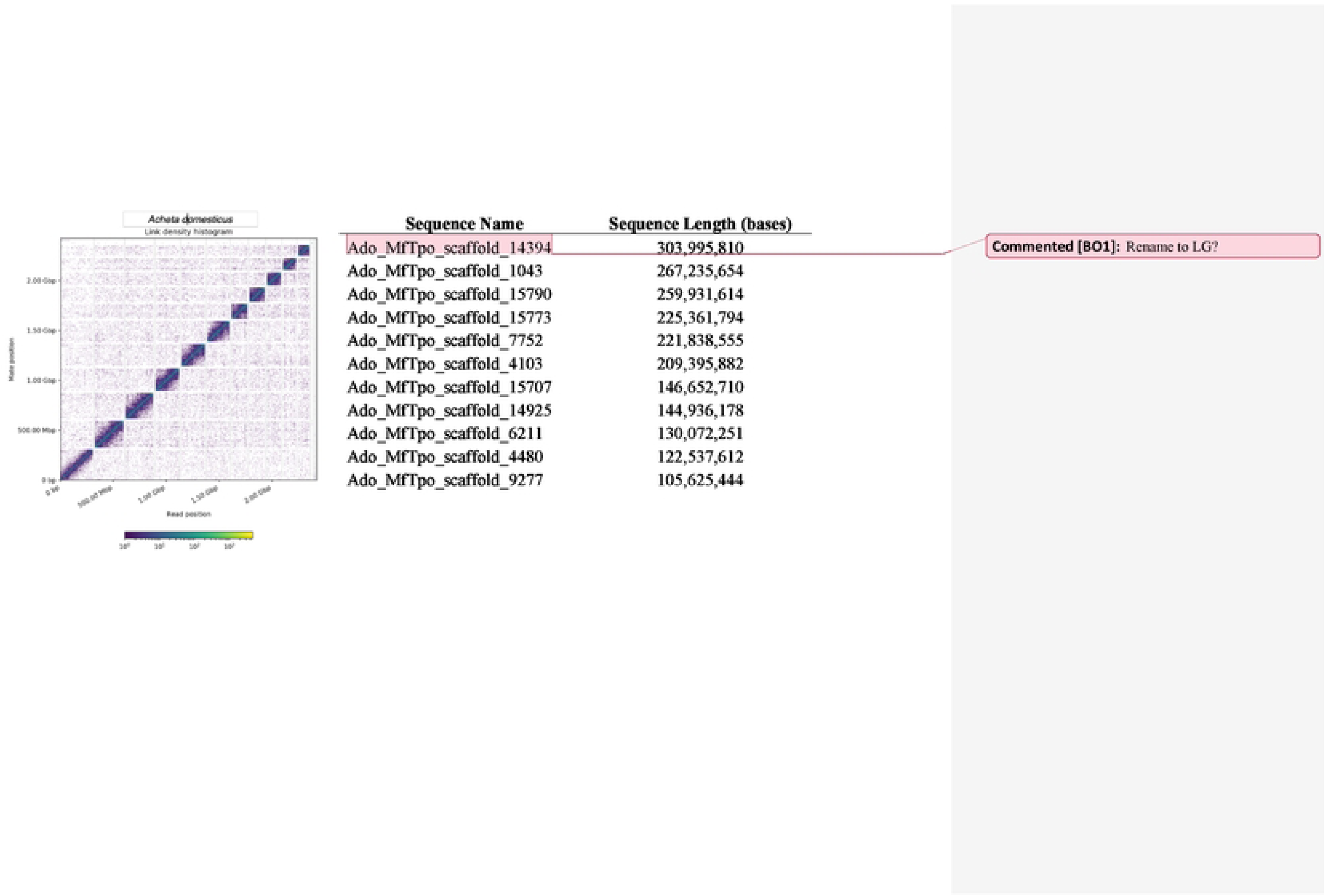
Link density histogram of the *A. domesticus* Hi-C scaffolded assembly (left) and scaffold name and length of the 11 largest scaffolds in the assembly (right).

**Table 1.** Metrics of the *A. domesticus* genome assembly, including the CANU draft assembly from PacBio long read data, and the scaffolded assembly from Chicago and Hi-C long range data.

| Metric | PacBio Draft Assembly | Chicago/Hi-C Scaffolded |
| --- | --- | --- |
| PB coverage | 27.3x (Sequel I) | - |
| #Contigs | 28,340 | - |
| Contig$N_{50}$ | 321,088 bp | - |
| Chicago/HiRise coverage | - | 10.37x |
| HiC coverage | - | 11,358.72x |
| Number of scaffolds | - | 16,290 |
| $N_{50}$ (Mb) | - | 221.839 Mb |
| Longest Scaffold | - | 303,995,810 bp |
| $L_{50}$ (#scaffolds) | - | 5 |
| Busco (Insecta) | - | 94.8% |

*In silico* genome annotation revealed 29,304 predicted genes in 1,064 scaffolds, concentrated in the 11 large scaffolds (84%). We compared the length of the BUSCO reference genes from *A. domesticus* to those from a model coleopteran genome, *T. castaneum* (Supplemental Figure S3). We found that 84% of the reference genes in *A. domesticus* were longer than those in *T. castaneum*, some more than 80-fold longer. In most case the longer length of *A. domesticus* genes was due to longer introns.

There were 6,246 scaffolds in the *A. domesticus* genome assembly that were identified as metagenome sequences using multiple automated and manual annotations. Metagenomic scaffolds were from 1,013 to 512,394 bp, and one larger scaffold, Ado_MfTpo_scaffold_8829, with 3,776,750 bases and 218 genes was identified as *Acinetobacter baumannii*; these scaffolds were submitted to NCBI as *A. domesticus* associated metagenome data. Approximately 83% of the metagenome scaffolds had similarity to Invertebrate Iridescent Virus 6 (IIV6, S3 File, “Ado_metagenome_Kraken2”). Bacterial scaffolds were mostly from the classes Gammaproteobacteria, Flavobacteriia, Sphingobacteriia, Fusobacteriia, Betaproteobacteria, Alphaproteobacteria, Actinomycetia, and Bacilli (Supplemental Figure S4b). However, follow-up screening of unclassified reads using blastn [70] and NCBInr found an additional 913 potential metagenome scaffolds (S3 File, “Blastn_unclassified”). The final scaffolds were identified as mostly IIV6 and bacteria, but also a smaller number of scaffolds potentially identified as fungi, protozoa, and nematode (S3 File, “Metagenome_summary”). Of note were 27 scaffolds from Wolbachia described from other insects. In our analysis we did not find evidence of common foodborne pathogens such as *Salmonella sp.*, and the few scaffolds aligning to *Listeria sp.*, *Escherichia coli* or *Staphylococcus sp.* were insufficient to confirm the presence of these pathogens.

A closer examination of the predicted proteins from the virus scaffolds indicated that only seven were assigned as “high-quality” viral genomes, and 25 scaffolds were determined to be viral genomes of “medium-quality” as determined by CheckV default parameters (Nayfach, et al., 2021; S4 file). Although many shorter scaffolds were not aligned to the IIV6 genome, many of the predicted genes in these scaffolds were annotated as viral. Therefore, these scaffolds are likely to be viral and quality may be affected by the database integrity, sequence read quality, or assembly artifacts. These sequences also may represent new viruses. Other sequences were classified as low-quality matches or undetermined genomes.

Interestingly, one medium quality scaffold (Scaffold_4233) was similar to a circular virus (DTR_035638) from the metagenome in the reduced digestive system of an oligochaete, *Olavius albidis*, a marine gutless worm related to another *Olavius* spp with a symbiont metaproteome that evidently contributes to nutrient metabolism [71]. This scaffold is longer (8,979 bp) than that of the circular virus (5,878 bp).

### Immunity genes

Conserved sequences involved in insect immunity were used to identify orthologss in the *A. domestica* genome assembly. Genes related to conserved cercropins and defensins were not found in the assembly and have not been identified in the literature from any orthopteran insect to date. However, there were two genes (S5 File) identified as peptidoglycan recognition protein (PGRP). Three transcripts, found in a previous transcriptome assembly [22], corresponded to these PGRP genes (Figure 2). Only one PGRP transcript appears to be expressed at low levels in the embryo. Expression of all PGRPs, especially ci_142522_1, ramps up during the 1^st^ week of development, and gradually decreases throughout the nymphal stages. PGRPs are expressed at higher levels in female compared to male adult *A. domesticus*.

**Fig 2.**
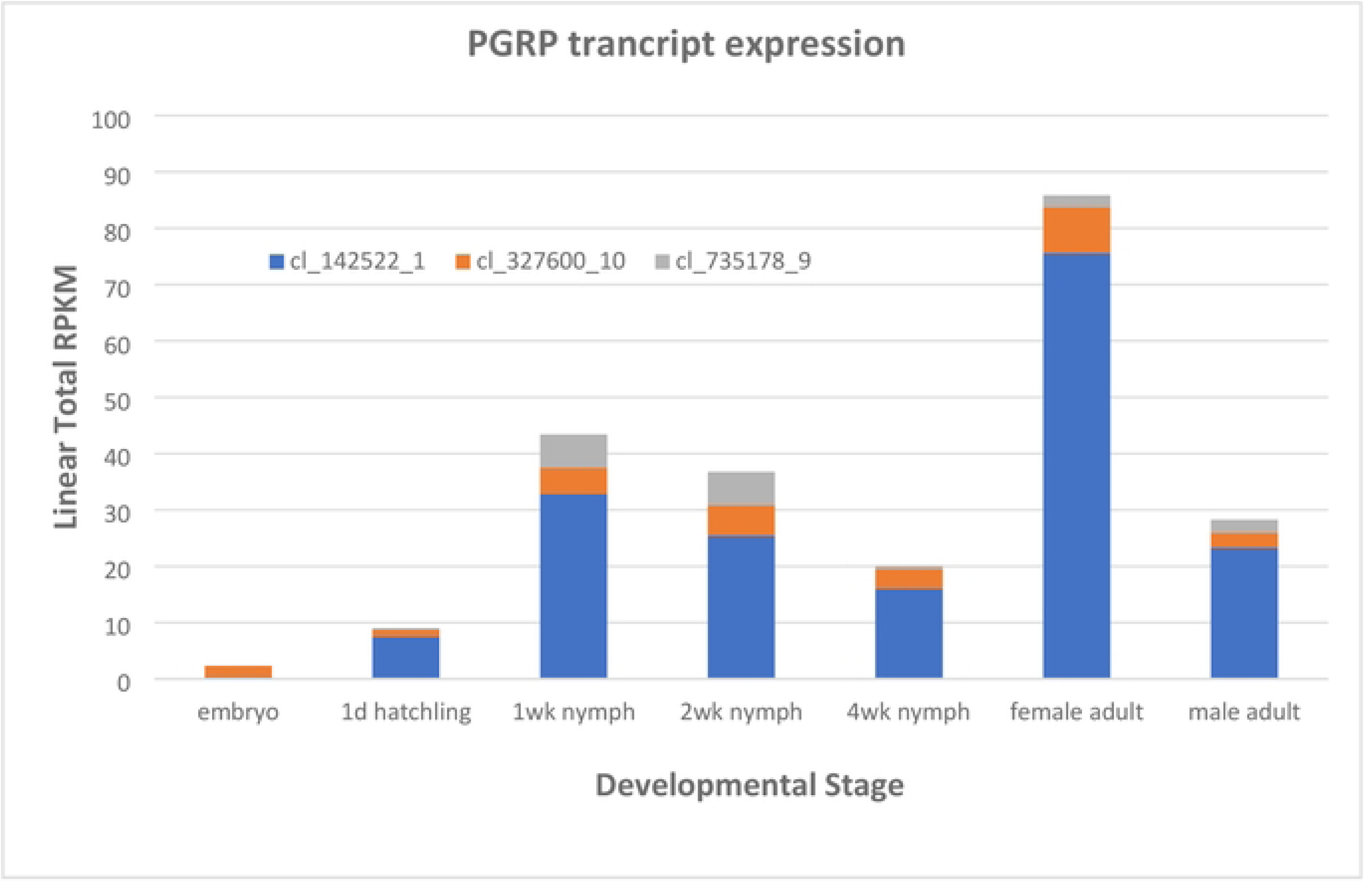
Expression profiles of three potential PGRP transcripts from *A. domesticus* developmental stages or male and female adults. The figure legend has the identification of transcripts in a previous transcriptome assembly [22].

Eleven genes were identified as gram negative binding proteins (GNBP, S5 File). These GNBP corresponded to 27 transcripts from the transcriptome assembly [22], some which were apparent isoforms of ANN05691-RA (5), ANN05692-RA (3), ANN05929-RA (4), and ANN16377-RA (3). Most of the GNBP transcripts were expressed at low levels in the developing *A. domesticus* adult (Figure 3A). However, cl_1105989_1 and _2 isoforms of ANN16377-RA were more highly expressed in all life stages except for embryos, as well as male and female adults.

**Figure 3.**
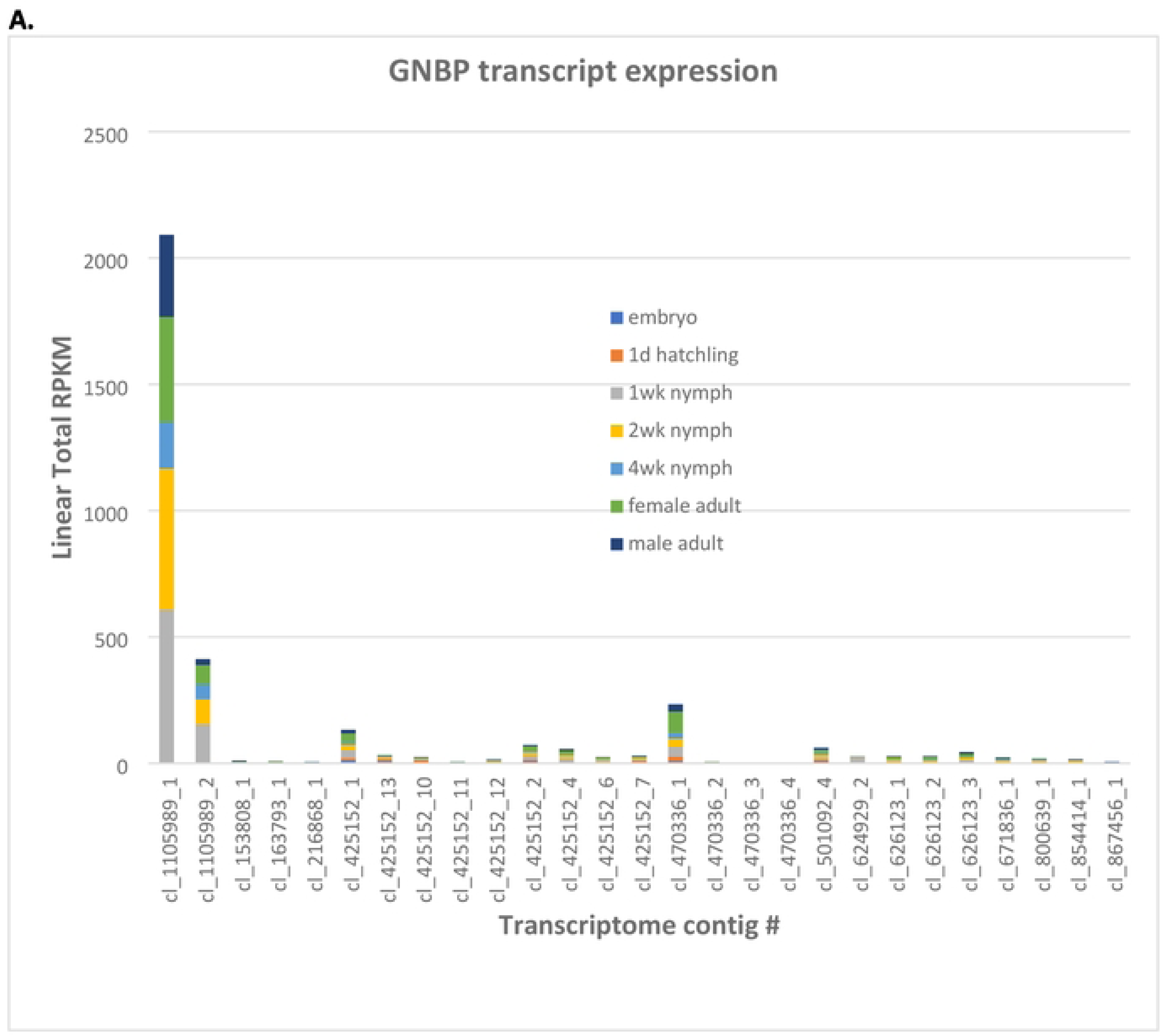
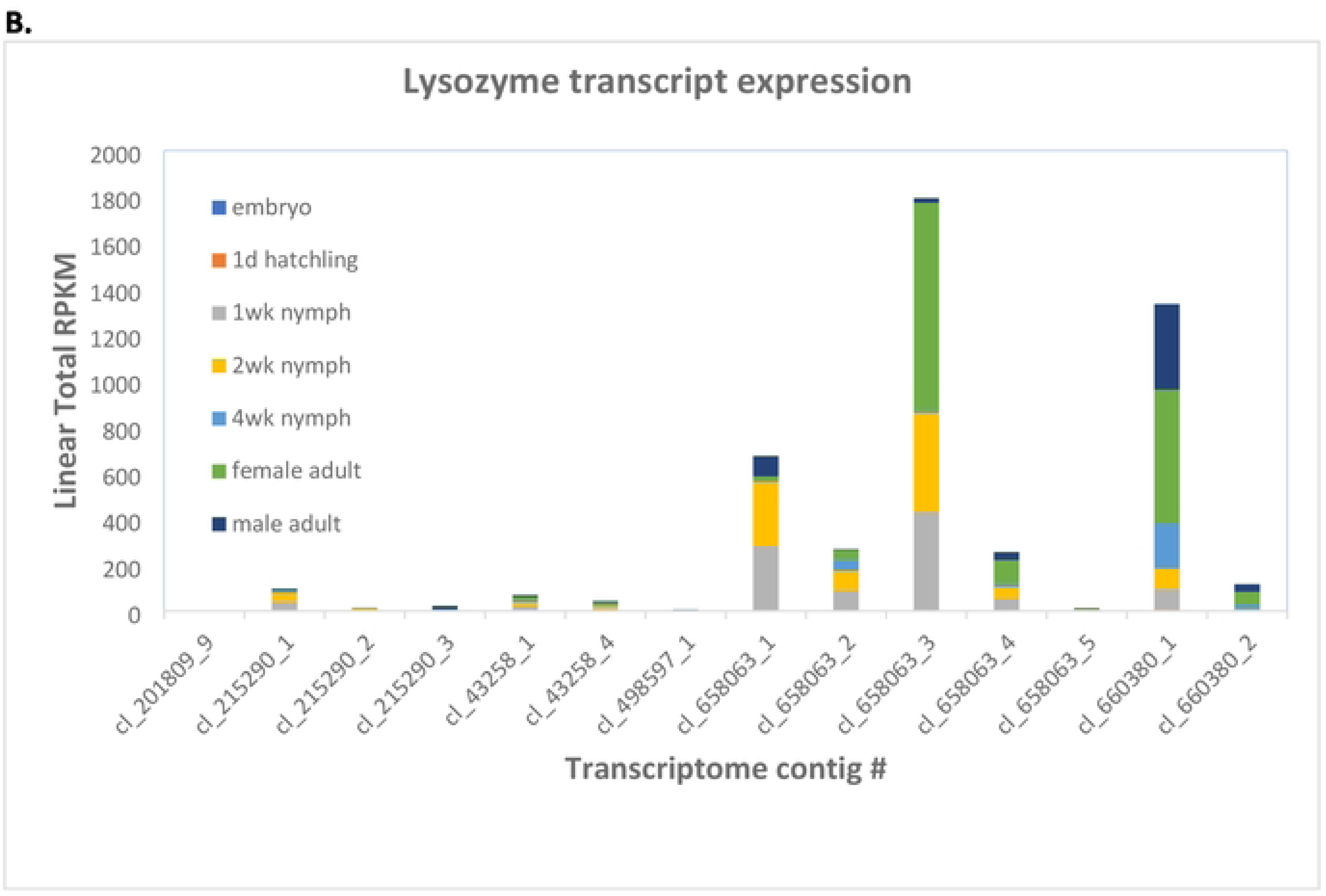
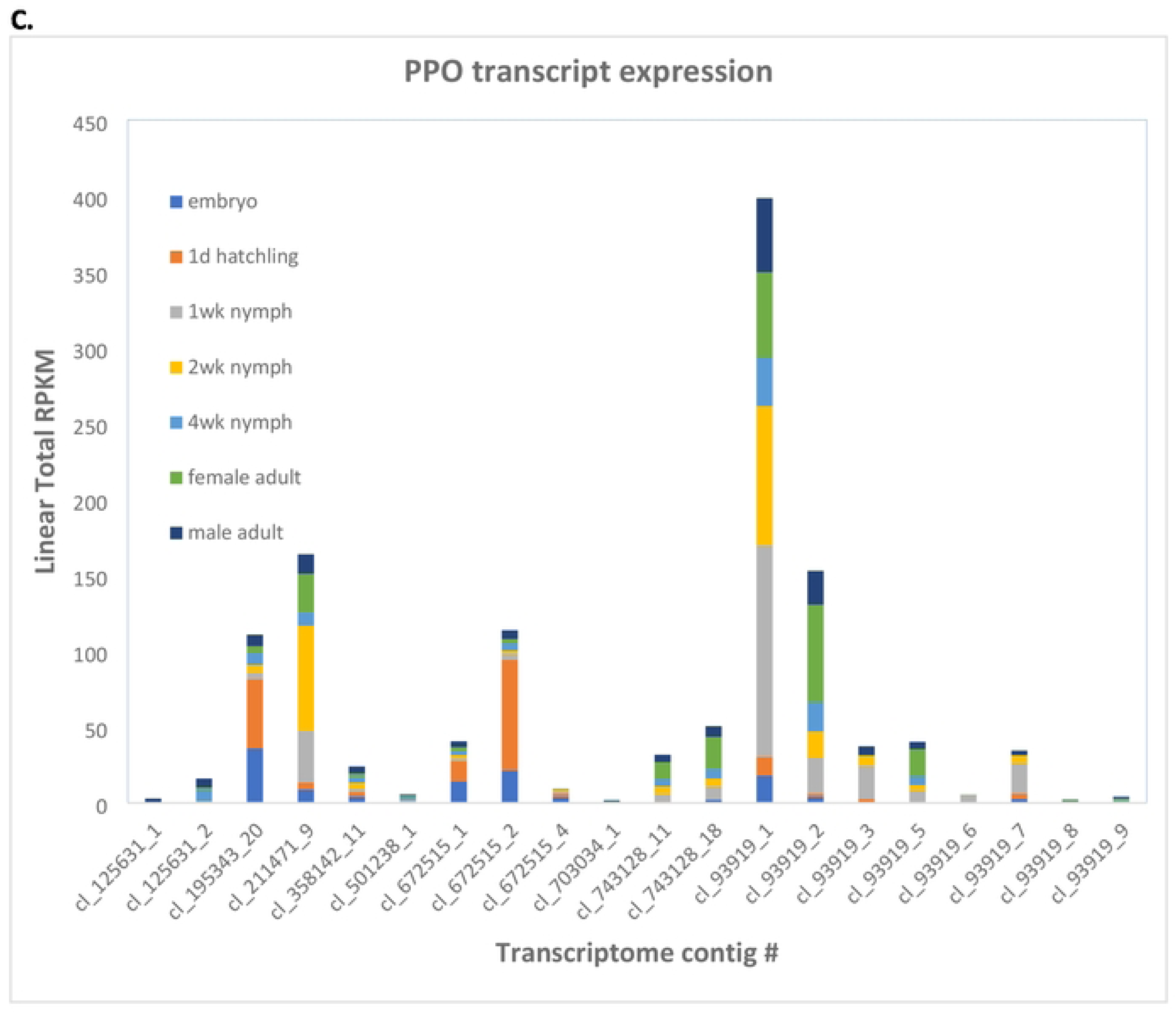
Expression profiles of transcripts tentatively identified as immune-related genes in *A. domesticus* developmental stages or male and female adults (as indicated in the legend). Transcripts were identified in a previous transcriptome assembly [22]. A. GNBP; B. Lysozymes; C. PPO.

There were four genes annotated as lysozymes in the *A. domesticus* assembly (S5 File). Contigs cl_658063 (five isoforms) and cl_660380_1 (two isoforms) corresponding to ANN24139-RA and ANN24140-RA/ANN24139-RA, respectively, were the most highly expressed putative lysozyme transcripts in the transcriptome assembly [22]; ANN19980-RA transcripts were expressed at low levels (Fig. 3B). Lysozyme transcripts were expressed higher in 1 and 2 wk nymphs, and male and female adults.

The *A. domesticus* genome assembly contained nine genes annotated as PPO (S5 File). Contig cl_93919 (nine isoforms), corresponding to all nine genes, was more highly expressed throughout life stages (Fig. 3C). However, cl_672515 (three isoforms), corresponding to ANN15897-RA, and cl_195343, corresponding to ANN15899-RA, were more highly expressed in newly hatched larvae.

### Repetitive Sequences

We annotated repetitive elements of the newly assembled genome of *A. domesticus* along with those in previously sequenced orthopteran genomes. We compared the repetitive elements in the genome assemblies of *A. domesticus* to those in other crickets (*Apteronemobius asahinai*, *Gryllus bimaculatus*, *Laupala kohalensis*, *Teleogryllus occipitalis*, *T. oceanicus*) and a locust, *Locusta migratoria*. This approach identified repetitive elements ranging from 33.00% of the *G. bimaculatus* genome to 58.13% of the *L. migratoria* genome (Table 2). Among the cricket genomes, *A. domesticus* had the highest overall content of repetitive elements at 49.42%. Among the major classes of the repetitive elements, DNA transposons were the most abundant element in the *A. domesticus* genome, accounting for 8.26% (Fig 4A). The major classes of repetitive elements in the cricket genomes examined were not uniformly present but were biased by species. For example, in the cricket genomes, *L. kohalensis* LINE had the highest abundance at 13.4%, while *G. bimaculatus* and *T. oceanicus* had only 3.34% and 4.04% LINE, respectively.

**Figure 4.**
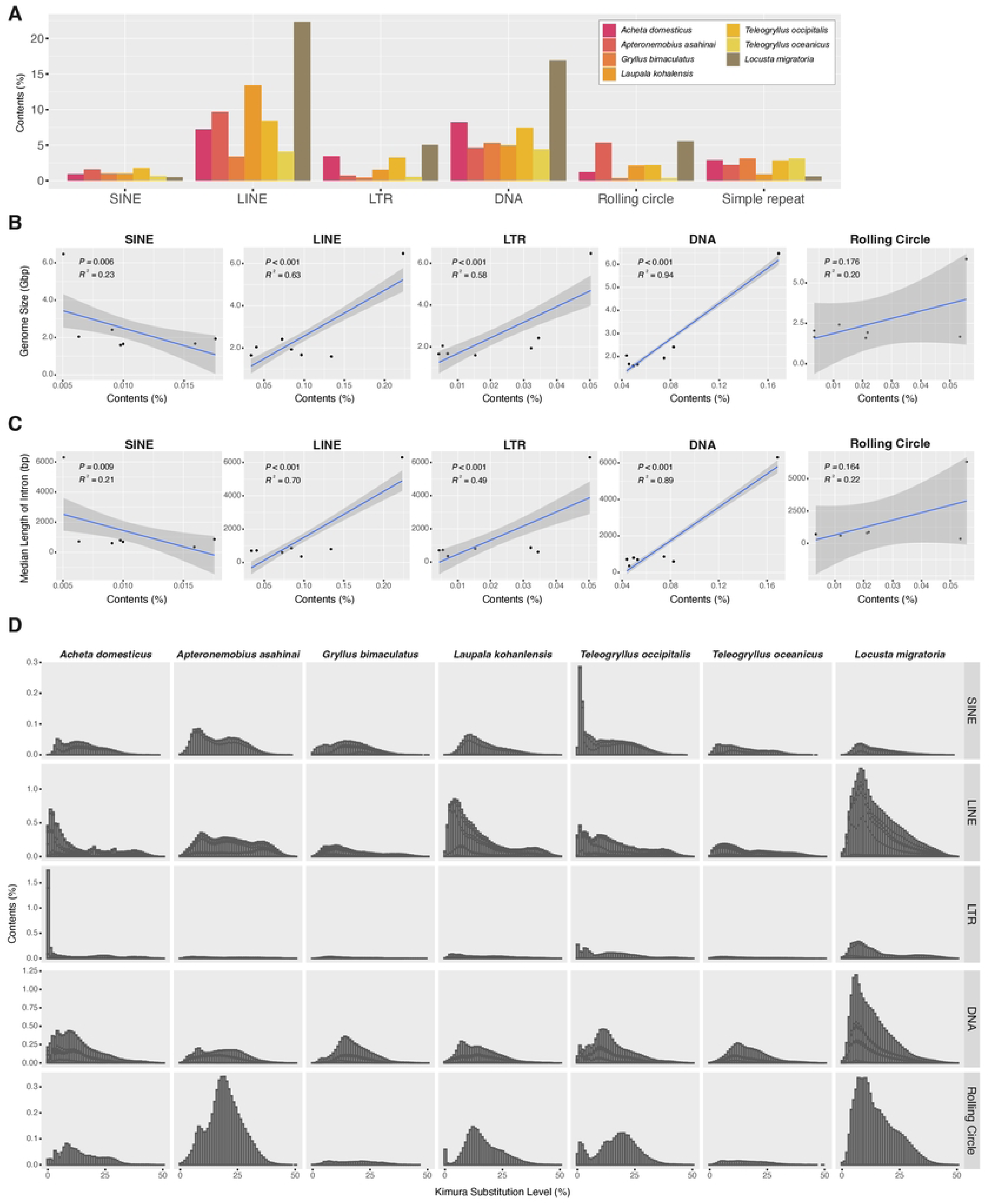
Transposable elements (TEs) in publicly available orthopteran genomes (six crickets, including *Acheta domesticus*, *Apteronemobius asahinai*, *Gryllus bimaculatus*, *Laupala kohalensis*, *Teleogryllus occipitalis*, *Teleogryllus oceanicus*, and one locust, *Locusta migratoria*). (A) Contents of representative repetitive elements including TEs in Orthoptera. The correlation between genome coverages of major TE classes (SINE, LINE, LTR, DNA and rolling circle) and genome sizes; (B) median length of intron; (C) Adjusted by Pearson correlation p-values (*P*) and coefficients (*R*^2^); (D) Repeat landscape of the major classes of TE classes (SINE, LINE, LTR, DNA and rolling circle) in the analyzed genomes. The Kimura substitution levels (x-axis) show sequence divergence, or “TE age”, for the major classes. The classes that are skewed to the left (less sequence divergence) are estimated to have more recently diversified histories than the classes that are skewed to the right.

**Table 2.** Comparison of the size of the genome of orthopteran species, repetitive content, and the mean length of introns.

| Species | Genome size (bp) | Repetitive content (%) | Mean length of intron (bp) |
| --- | --- | --- | --- |
| *Acheta domesticus* | 2,254,915,693 | 49.42 | 1,720.0 |
| *Apteronomobius asahinai* | 1,676,217,857 | 39.68 | 1,000.4 |
| *Gryllus bimaculatus* | 1,658,007,496 | 33.00 | 4,124.6 |
| *Laupala kohalensis* | 1,595,214,429 | 43.16 | 2,001.7 |
| *Locusta migratoria* | 6,476,667,755 | 58.13 | 11,988.4 |
| *Teleogryllus occipitalis* | 1,933,897,352 | 45.90 | 2,974.3 |
| *Teleogryllus oceanicus* | 2,045,067,382 | 38.91 | 2,453.7 |

Because repetitive elements, in particular transposable elements (TEs), are well-known contributors to genome size expansion in a wide range of insect species [72–73], we further examined the relative contribution of repetitive elements to the genome sizes of the orthopteran species. There was a significant correlation between total contents of repetitive elements and genome sizes in Orthoptera species (*P*=0.024 and *R*^2^= 0.61) (Figure S6a). Because the contribution of the large genome size of *L. migratoria* may be significant in this analysis, we excluded the data of *L. migratoria* in the analysis and found that there was no correlation (*P*=0.166 and *R*^2^= 0.27 (Figure S6b). We then dissected the repetitive elements into the major classes of TEs (i.e., SINE, LINE, LTR, DNA and rolling circle) and investigated their contributions to genome size. As a result, we found that LINE, LTR elements and DNA transposon may contribute to the expansion of genome size in Orthoptera (LINE, *P*<0.001 and *R*^2^= 0.63; LTR, *P*<0.001 and *R*^2^=0.58; DNA, *P*<0.001 and *R*^2^=0.94) (Figure 4B). When *L. migratoria* data was excluded, we found that the contents of LTR elements and DNA transposon were positively correlated with genome sizes (LTR, *P*<0.001 and *R*^2^=0.43; DNA, *P*<0.001 and *R*^2^=0.50) (Figure S6a).

We next studied associations among intron length, genome size, and repeat content in Orthoptera, including *A. domesticus*, because they have been reported to be mutually correlated [74]. Median length of the *A. domesticus* gene model was 602 bp, which was about one-tenth of that of *L. migratoria* (Table 2). We found that total contents of repetitive elements and median length of introns in orthopteran species, including locust, were positively correlated (*P*=0.044 and *R*^2^=0.51), but not within the cricket species (*P*=0.791 and *R*^2^<0.01) (Figure S6b). Further, correlations between intron length and major TE contents were positive for LINE, LTR elements and DNA transposon in Orthoptera (LINE, *P*<0.001 and *R*^2^= 0.70; LTR, *P*<0.001 and *R*^2^=0.49; DNA, *P*<0.001 and *R*^2^=0.89) (Figure 4C), but not for SINE and rolling circle. These correlations were no longer established except for DNA transposons after limiting the analysis to crickets only (DNA, *P*=0.034 and *R*^2^=0.05) (Figure S6b). There was also significant and strong positive correlation between the median length of introns and genome size in orthopteran species (*P*<0.001 and *R*^2^=0.97) (Figure S6c). However, when limited to the analysis within crickets, genome size did not correlate with intron length (*P*=0.949 and *R*^2^<0.01). Furthermore, a more detailed examination of the classes of TEs in Orthoptera revealed a positive correlation with intron length and genome size for LINE/CR1, LINE/L1, LINE/Penelope, LINE/RTE, DNA/hAT, DNA/Kolobok and DNA/TcMar (Table S7). These results suggest that repetitive elements, especially several classes of TEs such as LINE and DNA transposon, may contribute to variation in genome size and intron length in Orthoptera, and that they are unlikely highly correlated within closely related species belonging to the same family, but more likely highly correlated at the suborder level.

We also examined whether abundance pattern of TEs in orthopteran genomes is due to shared ancient proliferation events or recent/ongoing activity of TE. We analyzed the distribution of sequence divergence of the annotated TE to infer the timing of the change in TE composition. As a result, we found TE distribution patterns that were *A. domesticus*-specific or common to crickets (Fig 4D). In *A. domesticus*, distributions of LTR transposon, especially LTR/ERV1 and LTR/Gypsy, showed a high abundance of recently diverged TE sequences compared to the other species, indicating an ongoing proliferative activity (Figure S6d-f). Likewise, a similar pattern was seen in SINE of *T. occipitalis* genome. On the other hand, the distribution pattern of DNA transposons in the orthopteran genomes examined here is relatively similar among the crickets, suggesting a possible shared activity of DNA transposons in ancient lineages of the crickets.

### Promoters and genes annotated for genetic engineering

For our genetic engineering experiments, we needed to identify promoters of specific genes of interest. First, we identified and annotated the *vermilion* gene from *A. domesticus* (*Ad vermilion*) (Figure 5a) as our target for CRISPR editing and the muscle actin gene from *A. domesticus* (*Ad muscle actin*) (Figure 5b) for use of its promoter for gene of interest (GOI) expression. The *Ad vermilion* mRNA was identified from our previously published *A. domesticus* transcriptome data [22] by tBlastn using the *Tribolium castaneum vermilion* amino acid sequence [75]. An incomplete mRNA was identified as the *Ad vermilion* mRNA. We then used this mRNA sequence to find the *Ad vermilion* gene location in the genome. This allowed us to annotate the gene and map the intron-exon splicing sites to identify each exon. The *Ad muscle actin* gene and its promoter were identified the same way initially using *T. castaneum muscle actin* gene. In addition, we were able to identify two very similar actin genes in the *A. domesticus* genome. Since muscle actin genes in many insect species are a single exon gene, we were left with one likely muscle actin gene with the highest score from blast and having a single exon. Once the gene was identified, we searched genomic sequence 1.5 kb upstream from the prospective start codon for the possible promoter regions using Neural Network Promoter Prediction tool (Figure 5b).

**Figure 5.**
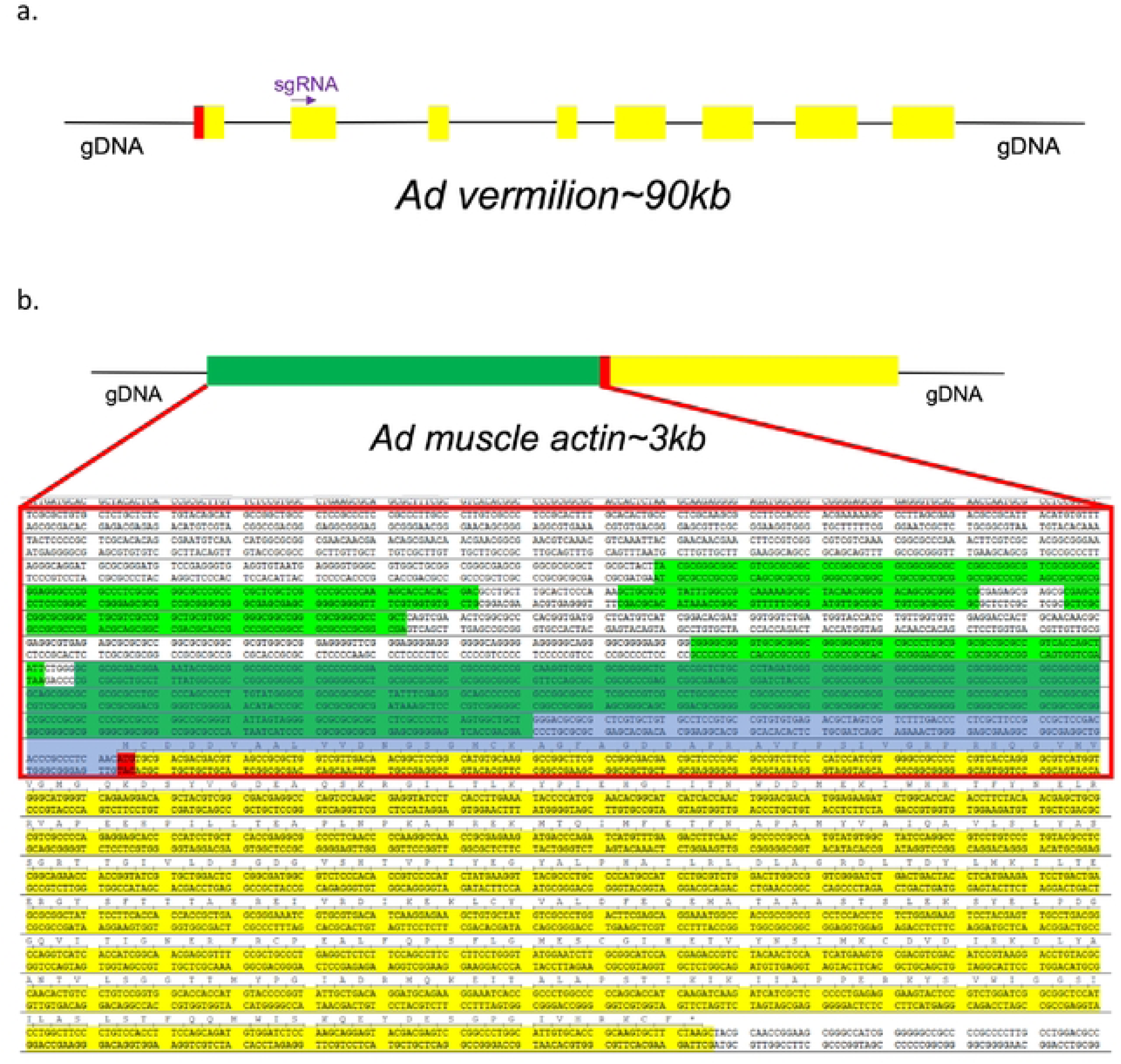
Design of CRISPR target in *A. domesticus*. a) Schematic of the annotated *Ad vermilion* gene from our *A. domesticus* genome data. Red box: ATG starting code; Yellow boxes: exons; Purple arrow: sgRNA site. Note: in this figure intron sizes are not exactly proportional to exons; b) *Ad muscle* actin gene with possible promoter areas. Red box: Sequences use for promoter prediction; Green sequences: predicted promoters; Red sequences: starting code; Yellow sequences: protein coding area; Gary blue area: promoter sequence use in CRISPR knock-in construct.

We targeted the *Ad vermilion* gene to also serve as a convenient visual marker (white/vermilion eye color phenotype) for preliminary identification of positive knock out / knock-in G^0^ crickets. The *Ad muscle actin* gene’s promoter was utilized due to its high level of continuous expression in all life stages as well as its characteristic muscle anatomy expression phenotype which is easily identified. For our downstream goals of expressing genes of interest in farm raised insects for food, feed and bioproduct manufacturing, it is usually critical to use promoters with high levels of GOI expression (eg: get the most product of interest per kilogram of insect) as well as have visual biomarkers (eg: white eye or EGFP phenotype being maintained through breeding selection) for simple genotype maintenance in farms and laboratories.

### CRISPR knock-in/out experiments

Having identified and annotated the *Ad vermilion* gene from *A. domesticus* (Figure 5a) we utilized the “GGP sgRNA designer” website tool to successfully identify three sgRNA sites as targets for our CRISPR knock-in/knock-out constructs (Table S8). The promoter of *Ad muscle actin* has many predicted promoters (Figure 5b). To keep the knock-in DNA construct size small, we used 455 bp of the possible promoter sequences upstream from the starting code as the promoter for the marker gene.

For knock-outs, we targeted *Ad vermilion* using CRISPR/Cas9. Using this target sequence, we designed and used either 1) a combination of 3 different sgRNAs or 2) a single sgRNA (sgRNA#1) (Table 3). The resulting G_0_ hatchling crickets which received all three sgRNAs had 68% eye color knock-out phenotype, while 32% of those receiving only sgRNA#1 had the phenotype. This phenotype resulted in eye color change from black (wild type) to vermilion or white eye color (Figure 6A,B). Our results show the knock-out color appears white in freshly hatched crickets and gradually changes to vermilion color after a few hours. In each set of injection experiments, both partially and complete knock-out of eye color were found (Figure 6B). Partial eye color knock out was mosaic. In comparison, the eye color in wild-type crickets were always black even in freshly hatched individuals (Figure 6A). To generate stable lines, groups of positive G_0_’s were subsequently self-crossed. For self-crosses, all 16 crosses and 9 out of 10 crosses from three and one sgRNA treatments provided vermilion eye color G_1_s. This indicates the CRISPR/Cas9 knock-out rate was particularly high in *A. domesticus*. To check the knock-out sequences, we tested five of the knock-out strains (G2) from the one sgRNA treatment. The sequencing results indicated deletion ranging from just a few base pairs to more than 240 bp among these five strains (Table 4).

**Figure 6.**
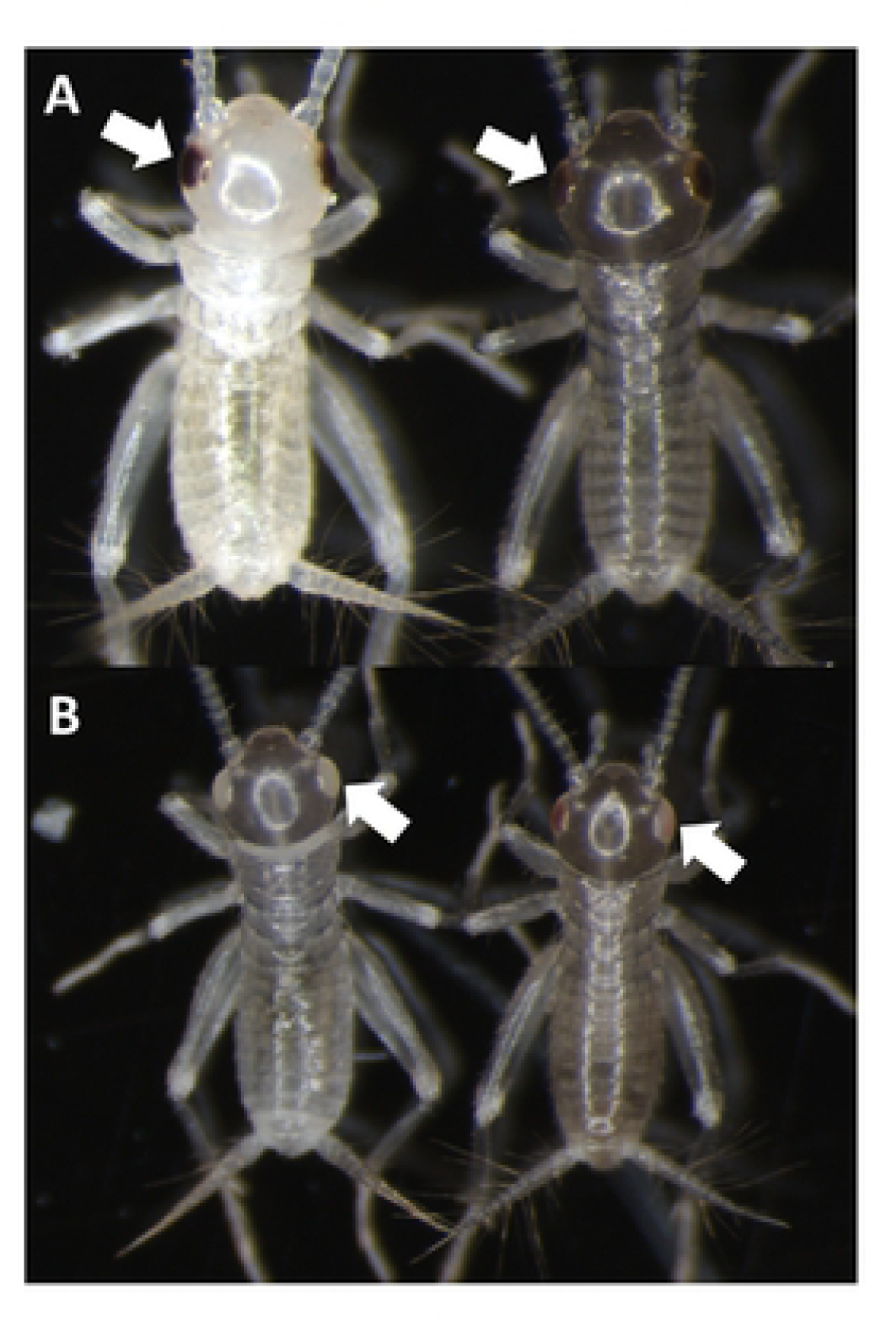
Vermilion eye color phenotype in cricket. A: wild-type eye color in freshly hatched cricket (left) and few hours old cricket (right). B: Eye color of completely knocked-out *Ad vermilion* gene in cricket (left) and cricket with partial knock-out phenotype (right).

**Table 3.** CRISPR knock-out microinjections and knock-out rate for *Ad vermilion* gene.

| sgRNA | Eggs injected | Hatched | Hatch rate | KO G<sub>0</sub>s* | # of crosses | KO G<sub>1</sub>** |
| --- | --- | --- | --- | --- | --- | --- |
| #1,2,3 | 1737 | 222 | 13% | 150 | 16 | 16 |
| #1 | 2065 | 433 | 21% | 139 | 10 | 9 |

\*Knock-out positive G<sub>0</sub>s
\*\* Number of crosses provide Knock-out positive G<sub>1</sub>s

**Table 4.**
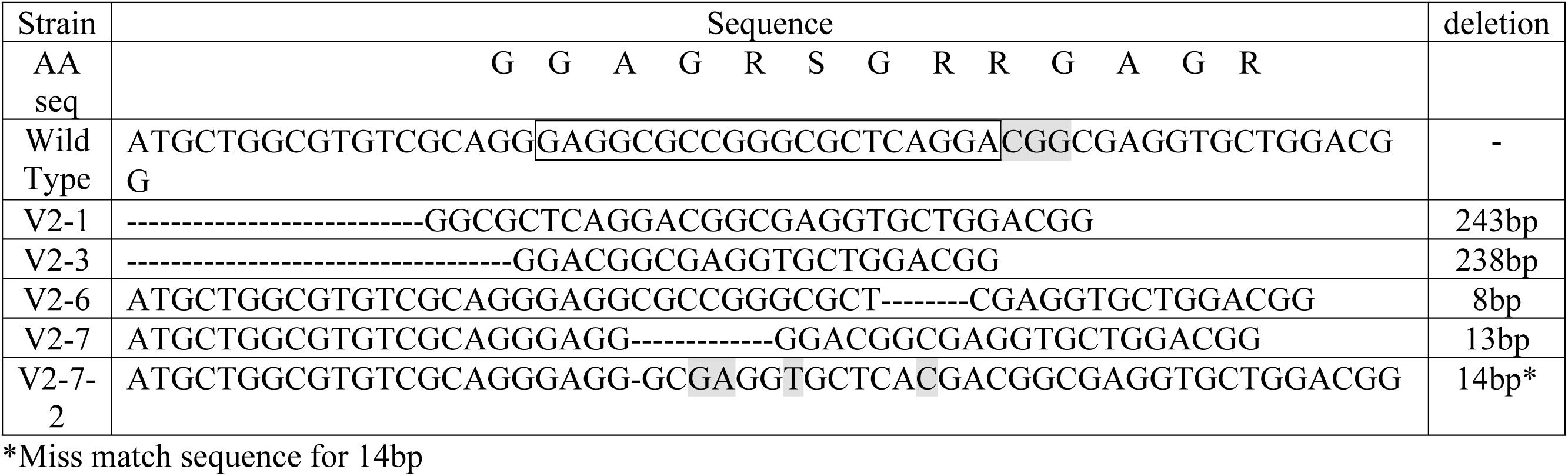
Deletions from CRISPR knock-out strains in house cricket.

For knock-in experiments, sgRNA#1 and the knock-in construct containing the enhanced green fluorescent protein (EGFP) marker gene coding region (Figure 7) were co-injected with Cas9 protein into young house cricket eggs. We injected 939 eggs and 255 crickets successfully hatched (hatch rate of 27%). From all hatched crickets, we obtained six G_0_ crickets showing EGFP expression (2%) (Figure 8). The knock-in positive phenotype selected was based on EGFP expression in muscle tissue due to use of the *Ad muscle actin* promoter. EGFP expression varied from a small group of muscles to more than half of the muscle in the body (Figure 8D,E). The enlarged picture shows a more detailed view of the muscle structure with GFP expression in part of the cricket leg (Figure 8F). These results indicate that the promoter is functional and successfully expressing the marker gene in the appropriate tissue. From the G_0_ EGFP positives, we were able to set up three out-crosses (with wild-type crickets) and one G_0_ EGFP negative cross (using injected hatchlings that did not show EGFP phenotype) to screen G_1_ offspring. From these experiments we received EGFP positive G_1_s from two of the positive G_0_ out-crosses (#1 and #3) and were able to establish two muscle EGFP expression colonies (Figure 9). In Figure 9A,B, we can see the EGFP expression patterns from #1 and #3 colonies are different. Crickets from colony #1 showed much lower EGFP expression and were only easy to screen from hind leg muscle, but most muscles in the body of crickets from colony #3 were expressing EGFP. There were no EGFP positive G_1_ offspring from the negative G_0_ crosses.

**Figure 7.**
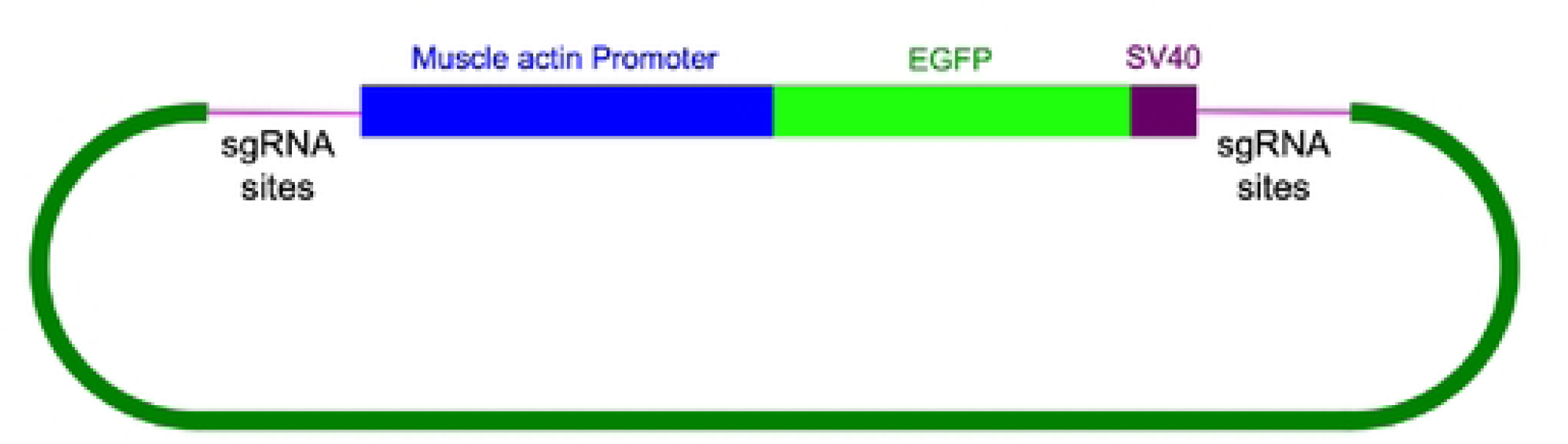
CRISPR knock-in construct. Using *Ad muscle actin* predict promoter driving EGFP as the marker gene. Two short sequences contain the *Ad vermilion* sgRNA sites are presenting on both side of the marker gene.

**Figure 8.**
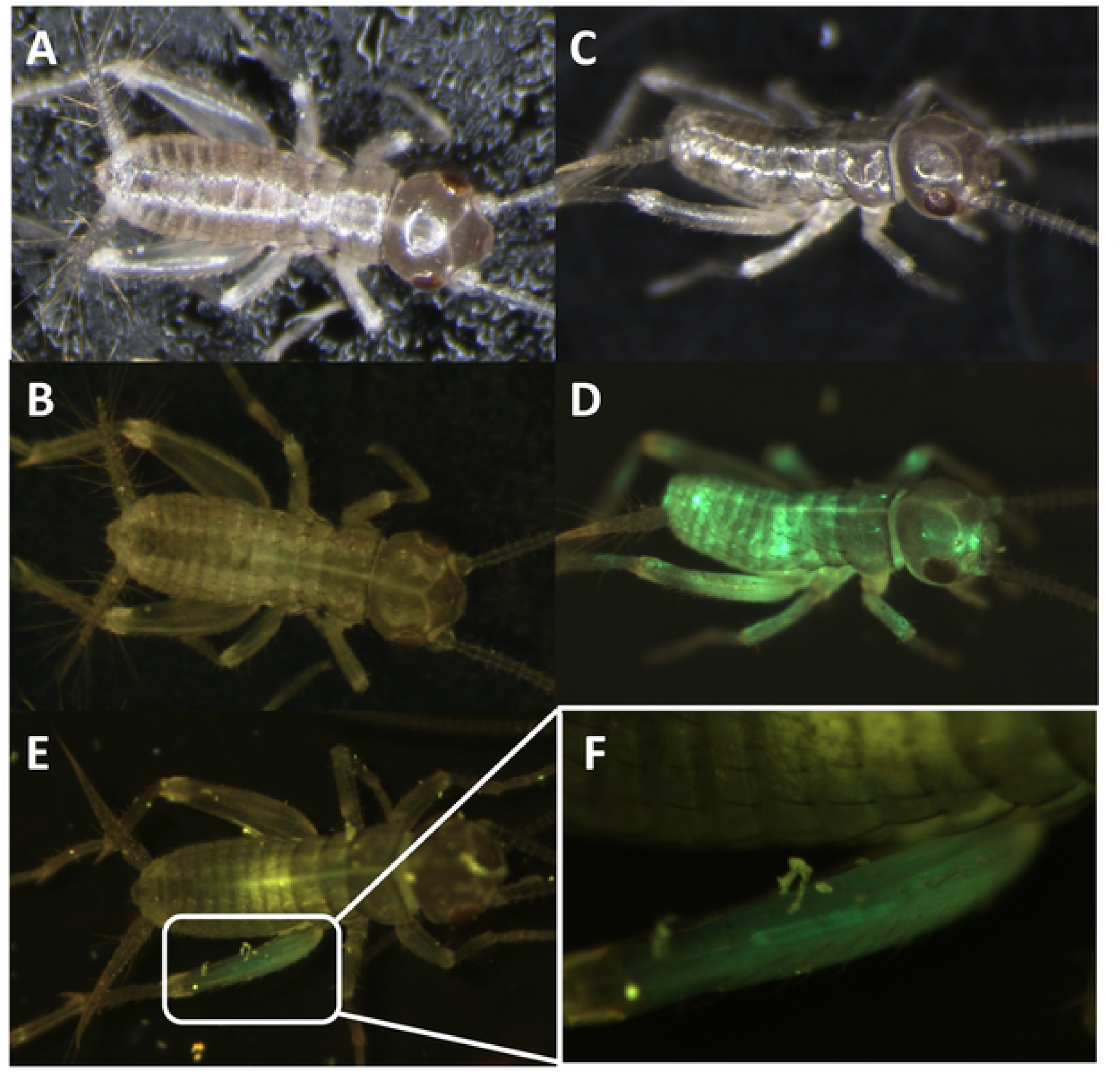
Somatic knock-in EGFP in cricket by CRISPR/Cas9. Wild-type cricket under white light (A) and florescent light with GFP filter (B). Successful knock-in cricket with large area of somatic knock-in under white light (C) and florescent light with GFP filter (D). Cricket with small area GFP knock-in phenotype (E) and enlarged picture of muscle structure GFP expression (F).

**Figure 9.**
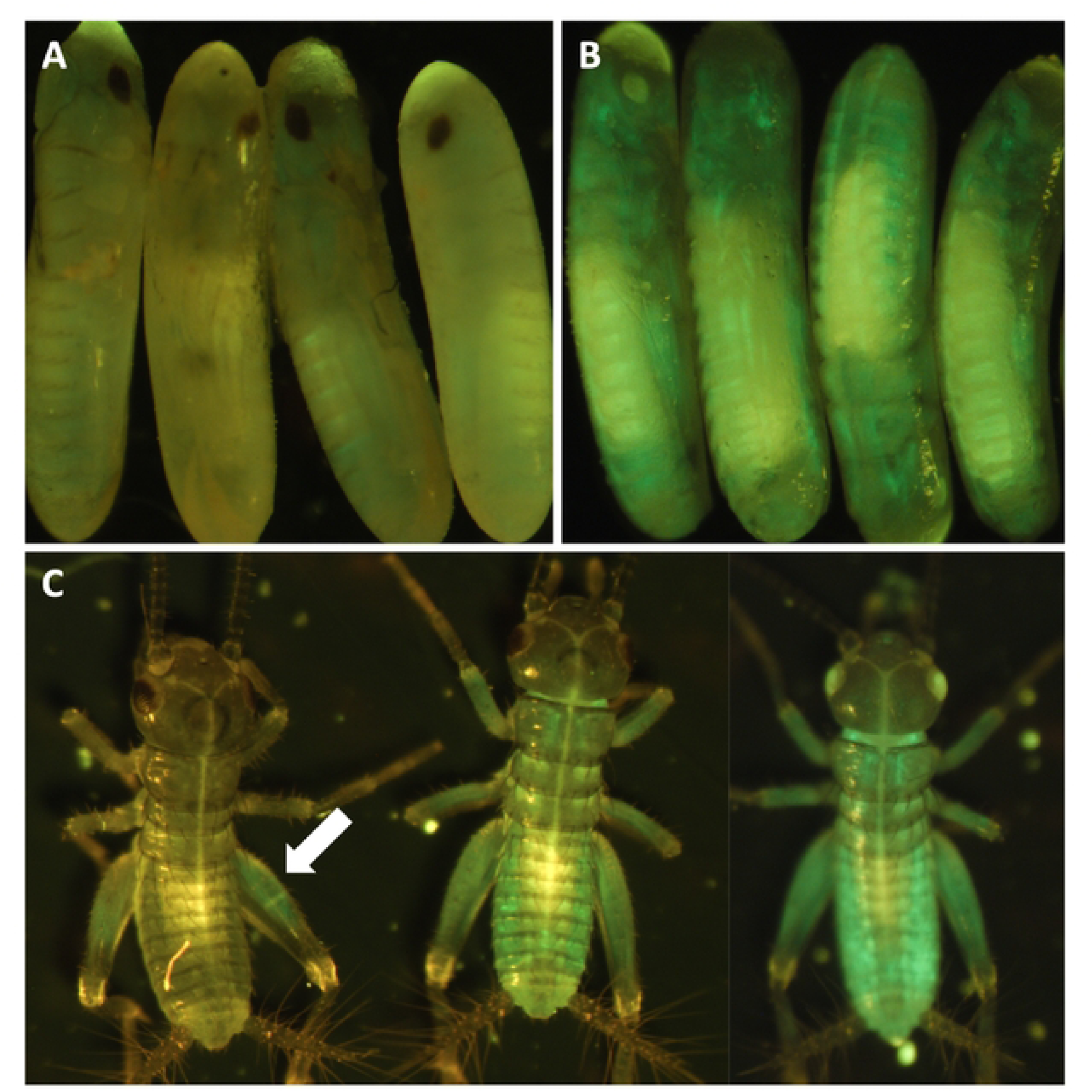
G_1_ crickets with GFP expression. (A) #1 G_1_ eggs with some GFP expression. (B) #3 G_2_ eggs with GFP expression. (C) GFP crickets from #1 (left) and #3 heterozygous and homozygous (middle and right) crickets. Note: Homozygous cricket with white eye phenotype.

### RNAi Knock-Down experiments

In addition to demonstrating knock-out and knock-in efficacy in house cricket based on our genomic data, we have also demonstrated efficacy of RNAi in this species using the same data. For this we used 2 different target genes to knock-out using different dsRNAs: 1) *Ad vermilion* gene (in wild type crickets) and 2) the EGFP gene we knocked in for our stable EGFP expressing lines. For each of the 2 respective cricket strains, we injected three sets of crickets for each dsRNA (Table 5). The results had survival rates (four weeks after injection) of approximately 70% for those with our RNAi construct and 51% for controls. Based on phenotype screening, either white eye or reduced EGFP expression, was approximately 86% (Table 5). For both phenotype screening, it took at least two weeks after injection before the RNAi phenotype was visible. The eye color knock-down phenotype was not as strong as the CRISPR knock-out, but it was sufficient to identify the difference (Figure 10A). The GFP RNAi knock down experiments showed similar result which can be identified starting at two weeks after injection (Figure 10B). We continued screening both phenotypes for five weeks after injection and noticed that both eye color pigmentation and GFP expression continued to be reduced between two and four weeks after injection.

**Figure 10.**
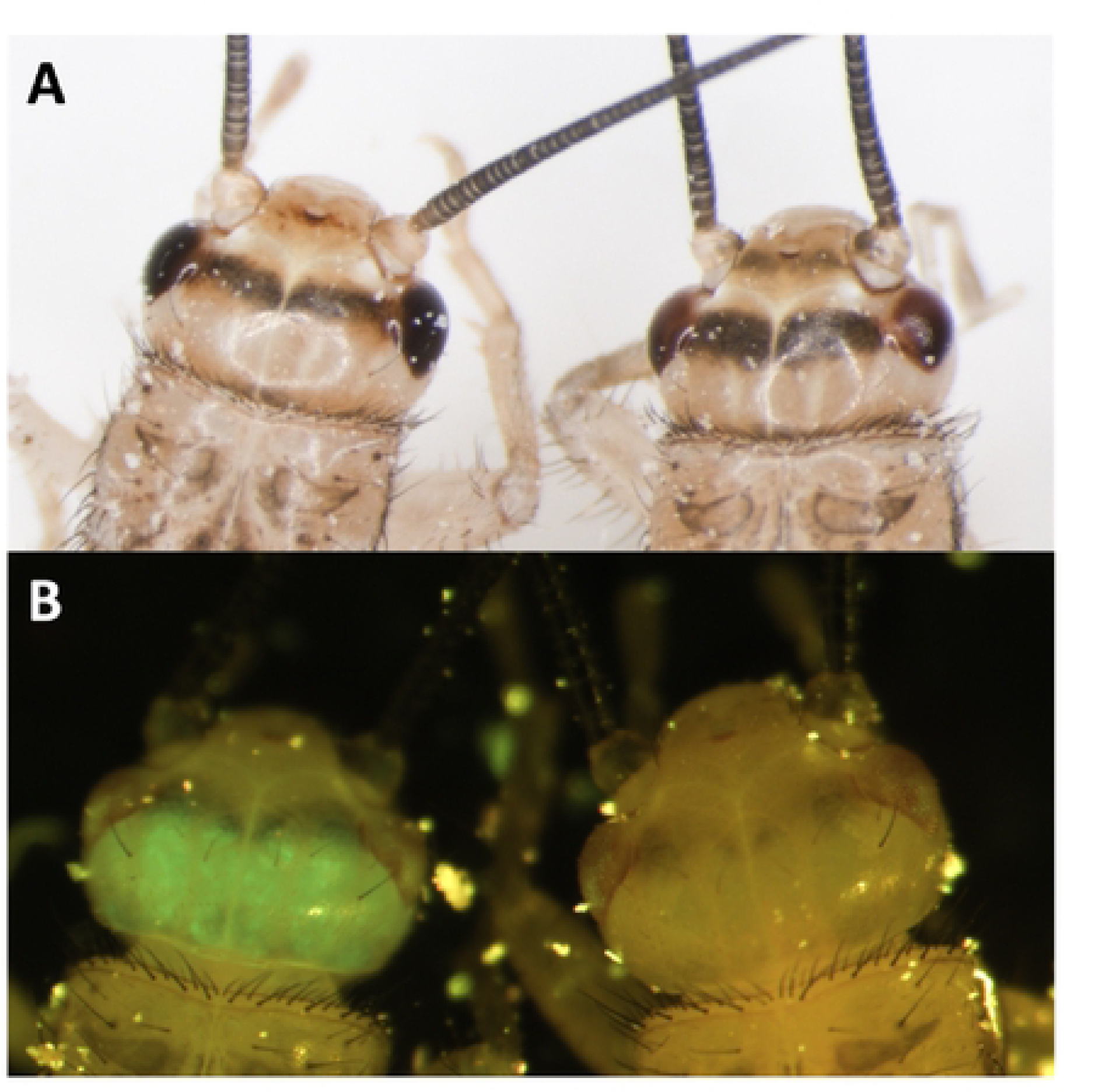
Cricket RNAi phenotypes. (A) *Ad vermilion* gene knock-down by RNAi in house cricket. Wild type eye color (left) compared with RNAi eye color (right). (B) GFP knock-down by RNAi in our EGFP expressing house cricket strain, AdVELV1-3. Regular EGFP expression (left) compared with dsEGFP RNAi cricket (right).

**Table 5.** RNAi results in *A. domesticus*.

| Name | Injected | Survival* | Survival rate | KD | KD rate |
| --- | --- | --- | --- | --- | --- |
| dsAdV-WT #1 | 19 | 16 | 84% | 13 | 81% |
| dsAdV-WT #2 | 19 | 17 | 89% | 13 | 76% |
| dsAdV-WT #3 | 20 | 10 | 50% | 8 | 80% |
| dsEGFP-WT #1 | 25 | 13 | 52% | 0 | 0% |
| dsEGFP-WT #2 | 18 | 9 | 50% | 0 | 0% |
| dsEGFP-AdVELV1-3 #1 | 20 | 12 | 60% | 12 | 100% |
| dsEGFP-AdVELV1-3 #2 | 20 | 17 | 85% | 15 | 88% |
| dsEGFP-AdVELV1-3 #2 | 17 | 9 | 53% | 9 | 100% |

\*Survival rate is count at four weeks post microinjection.

## Discussion

### Genome

The *A. domesticus* genome assembly represents a contiguous assembly for downstream applications. The assembly contains 11 large scaffolds, presumably representing most of the autosomal and X chromosomes, in 2.138 Gb, close to the flow-cytometry predicted genome size of 2.150 Gb for male *A. domesticus*. There were 29,304 predicted genes, mostly concentrated on the large chromosomes.

Immediately after obtaining the genome assembly, we looked for promoters for genes for use in our transgenic experiments but encountered genes with very long sequences that were distinctly longer than the coleopteran genomes we have studied. A quick comparison of the length of the BUSCO reference gene sequences estimated that the *A. domesticus* genes were on average 80% longer than those in *T castaneum.* We also noticed that, in most cases, the longer genes were due to longer introns that may be due to TEs as has been observed in Orthoptera [72–73]. Therefore, we compared repetitive elements in the *A. domesticus* genome assembly with those available for other orthopterans and found that the *A. domesticus* genome assembly contains the highest content of TEs (almost half of the assembly) compared to other cricket genomes. Repetitive elements and genome size in Orthoperta were significantly correlated if we included the very large genome of *L. migratoria* (*P*=0.024). When *L. migratoria* data was excluded, LTR elements and DNA transposon were positively correlated to the expansion of genome size (*P*<0.001). As we suspected, repetitive element and median length of intron were positively correlated (*P*=0.044) but only when *L. migratoria* was excluded, and especially for elements LINE/CR1, LINE/L1, LINE/Penelope, LINE/RTE, DNA/hAT, DNA/Kolobok and DNA/TcMar. Our data suggest that LINE and DNA TEs contribute to variation in genome size and corresponding intron length in Orthoptera, but this is likely found only at the suborder level. Furthermore, the TE distribution pattern of LTR transposons shows a recent or ongoing burst in the *A. domesticus* genome and likely contribute to the high abundance of repetitive elements in the genome of this species.

In addition to being a food source for animals and humans, crickets have also been utilized as a valuable model system for studying biological processes such as neurobiology, developmental biology, animal behavior and others [76]. With the recent interest in industrial farming crickets for human consumption, even greater potential exists for studying these creatures via expanded access to all life stages from farms, as well as the potential impact of research on cricket biology and genetics.

### *A. domesticus* Metagenome

Of the 16,290 original scaffolds in the *A. domesticus* assembly, 6,246 scaffolds were removed and submitted to NCBI as metagenome data. Most of these metagenome scaffolds had similarity to IIV6, but we were unable to assemble these scaffolds into a complete IIV6 sequence. Closer examination of the predicted proteins from the virus scaffolds indicated that only 32 scaffolds were high-medium quality viral genome sequences. Therefore, scaffolds with similarity to virus sequences at the nucleotide level may be sequencing artifacts, or some may represent novel viruses that have not been described in current databases. Importantly our analysis did not identify common foodborne pathogens such as *Salmonella sp*., *Listeria sp.*, *E. coli* or *Staphylococcus sp*.

The question of crickets harboring latent iridovirus infections was recently addressed [77]. In that study, iridovirus sequences were found in sick and healthy insects, and suggest that difference in immunity may prevent active diseases in healthy colonies. Our insects came from an insect farm with no reports of insect disease. We did find scaffolds similar to *Wolbachia* symbionts in other insects. *Wolbachia* has been reported to have a protective effect against viruses in some insects [78]. Further, Wolbachia was reported in a previous study of microbial communities in *A. domesticus* [79]. More research is needed to understand the prevalence of iridoviruses in farmed insects as well as the mechanisms that prevent active disease in infected but healthy individuals. It may even be possible to engineer virus resistant crickets through CRISPR genetic engineering technology. This is an important reason we seek to understand genes related to immunity in these insects.

### Immunity-Related Genes and Antimicrobial Peptides

We annotated genes related to immune responses in *A. domesticus* to better understand how these crickets respond to infection, important in insects being reared in close confines for animal feed and human consumption. The current study did not identify analogues of classical canonical small insect antimicrobial peptides (AMPs), such as cecropins or defensins, in the *A. domesticus* genome. Insects depend almost exclusively on an innate peptide and antimicrobial secondary metabolite forms of immunity against infections/pathogens [18, 82]. AMPs are found more in holometabolous than hemimetabolous species [80]), perhaps due to sampling (i.e., originally identified in insect pupae/larvae). However, this may also be due to the poor mobility and frequent terrestrial lifestyle of insect larvae vs nymphs (eg: grubs/caterpillars vs cricket/grasshopper nymphs), thus exposing holometabolous insects to be a wider variety of microbial pathogens for longer periods of their life [80]. Insect immune-related proteins have activity against microbes including Plasmodium sp. [83]. However, as with much of Class Insecta, insufficient research is available to identify biological activities and potential practical applications for these proteins [18]. Given their abundance in insects and the growing scale of the insect production industry, insect antibacterial, antifungal, antiparasitic, and antiviral proteins also could be a valuable and low-cost bioresource or byproduct for future biomedical applications. Many studies have identified numerous AMPs from insects with a wide variety of efficacies against various pathogens and other potentially valuable biological activities [84–85]. Thus, these proteins represent an untapped resource for potential therapeutic applications.

Instead of AMPs, we found other conserved genes in the *A. domesticus* genome assembly that may protect *A. domesticus* from pathogens, including PGRP, GNBP, lysozymes, and phenoloxidase. We reverted to our previous transcriptome data [22] to understand how these genes are expressed in different life stages or in male vs female adults. Two PGRP genes are increasingly expressed during early nymphal development and are more highly expressed in female than male *A. domesticus.* Eleven genes were annotated as GNBP, but only one gene, ANN16377-RA, was more highly expressed in nymphal and adults. Lysozymes and PPO are important in resistance to disease in crickets [81]. Overall, lysozyme genes were expressed higher in 1 and 2 wk nymphs, and male and female adults, PPO genes were expressed mostly throughout life stages, but two genes, ANN15897-RA and ANN15899-RA, were more highly expressed in newly hatched larvae and thus at a time the cricket would be upregulating immune defenses. These data will be important in developing protocols for healthy farm-reared insects, both in diagnostics and prophylatics for disease control.

Additionally, when considering the impact of insect genetic engineering such as work derived from this publication, insects may be an ideal host for expression and mass production of exogenous antimicrobial peptides from various species (eg: crickets or mealworms may be useful as bioreactors to produce a particularly active/valuable peptide from something like a moth which may be more challenging or costly to mass produce). As such applications are identified, given the scale of insect biological diversity as well as having an established insect mass production industry at the scale of the current food industry will allow a massive and low-cost resource for purifying these compounds for use in those applications.

### Genetic Engineering and Gene Editing via CRISPR

The primary purpose of this work was to sequence the *A. domesticus* genome to obtain necessary sequence data for genetic engineering to improve these animals as a sustainable food crop and for other forms of bioproduction. Thus, we identified and annotated two genes, *Ad vermilion* and *Ad muscle actin*, which were used to create knock-out ad knock-in strains of crickets as an initial proof of concept to develop the tools required for our downstream technologies. The analysis of *Ad vermilion* was our first indication that the genome of this species contains quite large introns. The exon sizes were similar to those of other orthologous insect genes such as *Tm vermilion* from *T. molitor*, but due to the intron extended length, this gene is more than 100 times longer in *A. domesticus*. The *T. molitor vermilion* gene is around 2 kb but is around 90 kb in *Ad vermilion*. On the other hand, annotation of the *Ad muscle actin* gene was more straightforward. As with *Tribolium sp.* and *Drosophila sp.*, *Ad muscle actin* has only one exon, and therefore is similar in size to that of other insect species. Once we had these genes annotated, we were then able to use the *Ad vermilion* gene to identify CRISPR target sites and the promoter region from *Ad muscle actin* gene for high level expression of knock-in marker genes. The *Ad vermilion* gene, when knocked-out and/or interrupted as a result of being a target for knock-in experiments, also functions as a convenient biomarker, as reducing or eliminating function of this gene results in a white-eye (or vermilion color eye) phenotype with less eye pigmentation than wild type. Thus, for the CRISPR genetic engineering work, we showed that the CRISPR/Cas9 gene editing tool works in *A. domesticus*. With up to a 68% CRISPR knock-out rate in G_0_, the efficiency is sufficient to produce and establish the modified strains for research and future commercial applications. After multiple generations in the laboratory, the *vermilion* mutation cricket colonies do not show any obvious decrease or increase in fitness based on survival and growth rate (data not shown). For our knock-in CRISPR experiments, we used the same sgRNA target site as for the knock-outs. As such, we expected the Cas9-sgRNA created DNA double break in both the insect genome and the knock-in plasmid. Therefore, the EGFP marker gene was likely incorporated into the genome through the non-homologues end joining (NHEJ) DNA repair process. We have shown that gene knock-in using CRISPR/Cas9 gene editing works well in *A. domesticus*. The *Ad muscle actin* promoter was successfully utilized to express exogenous EGFP as expected in muscle tissue. The G_0_ somatic knock-in rate was around 2%, and two out of three outcrosses provided EGFP positive G_1_, indicating the knock-in is efficient in those G_0_. As this was our first attempt, an optimized protocol will be created in the future to further improve this knock-in rate. Interestingly, the EGFP colonies we established showed different muscle EGFP expression patterns. This could be due to a number of reasons, such as the promoter was damaged during knock-in since we used NHEJ to create the knock-in, or due to epigenetic factors. Based on these results, we successfully showed CRISPR gene editing works to knock-in genes and function in house cricket. This groundbreaking experiment opens up many possibilities for valuable applications going forward which are discussed below.

### RNAi in A. domesticus

For the RNAi experiments, abdominal microinjections of young cricket with our target gene dsRNAs resulted in either eye color reduced phenotype or whole body GFP knock-down, depending on the construct and cricket strain injected. This indicates *A. domesticus* has the necessary genes for systematic RNAi and that they are functional, as was previously described in this insect [86] and in the *Allonemobius socius* complex of crickets [87]. From experience with other insect species, we expected RNAi to start showing phenotypic changes around 48 hours after injection or within 3-5 days at the earliest. However, in our case, since crickets regain their eye color every time they molt, we did not see the eye color change until their next molt after dsRNA microinjections. Depending on the timing, the eye color was sometimes more reduced after next two to three molts. However, since RNAi could only knock down the existing gene expression, it is likely eye color will gradually be restored through time within each molt. Thus, the RNAi eye color phenotype was as obvious as CRISPR knock-out. EGFP knock-down also showed visible phenotype (reduced EGFP expression under fluorescent microscope) after two weeks. We believe this slow response compared with other insects may be because the EGFP protein is more stable, so it takes longer for the pre-experiment protein to be degraded. For these RNAi experiments, we observed a lower survival rate in some of the microinjection groups likely due to non-optimal rearing conditions such as utilization of small containers for observation which were not large enough for more than 2 molts/live stage increases, which can be optimized in the future RNAi experiments.

This work provides the foundation for the potential of genome editing applications in farm raised house crickets such as disease resistance and alteration of the nutrient profile for production of numerous bioproducts using the low cost, low-tech farm raised insect bioreactor system. Insects are already highly efficient and sustainable compared with other livestock and protein production systems, as discussed above. Producing insects with higher growth rate, lower mortality due to disease and other factors and that are more nutrient dense through gene editing technology such as presented here could be a game changer to make them orders of magnitude more efficient and sustainable. What’s more, insects hold promise for other uses beyond a food/feed source. Many opportunities exist for insects as an untapped resource for applications such as pharmaceuticals including antimicrobials and antiviral substances, drug lead compounds, material for bioprospecting, enzymes, bioactive peptides, oils, biocompatable materials for wound healing and other medical applications, food waste mitigation, nutrient cycling, biodegradable plastics and packaging, new materials from chitin with novel properties and durability and many other applications [18, 37]). However, very little bioprospecting of insects has been done to date compared with other taxa. Beyond the substances that insects contain naturally, genetic engineering opens additional potential for efficient, low-cost bioproduction of vaccines, enzymes, antibiotics, antimicrobial peptides, antibodies, color pigments and dyes, flavors, fragrances, functional ingredients, and many others.

Developing the tools for insect genetic engineering, including high quality genomes, provides an open-ended opportunity to use insects for purposes besides mere sources of food, protein and dietary nutrients. We believe this foundational research will play a critical role in reducing human environmental impact by utilizing the largest, most diverse yet almost entirely untapped biological resource on Earth: Class Insecta.

## Materials and Methods

### House cricket colony maintenance

*A domesticus* were originally purchased from a US insect farm in 2018 and maintained as a lab colony for multiple generations. They were fed a diet of specially formulated cricket feed from LongStar (TFP Nutrition) (Nacogdoches, TX, USA), which was the same feed used by the insect farm. Feed and water were provided to each cricket cage twice per week. Eggs were collected twice per week from wild type adults in a petri dish (150 x 10 mm) filled with hydrated (deionized water saturated) polyacrylamide water crystals (these crickets prefer to lay eggs in moist areas). Each week, we replaced the old egg-lay dish with a new dish on Monday and Friday. Eggs were collected each Tuesday for colony maintenance. For egg collection, the petri dish with water crystals containing eggs was washed into a glass beaker using deionized water. Additional water was added to the beaker and stirred a few times, and eggs were allowed to settle to the bottom for approximately 10-15 seconds. Most of the excess water and crystals were slowly removed to retain eggs. Washing was repeated 2-3 times and a plastic Pasteur pipette was used to aspirate eggs from the bottom of the beaker and transfer to a piece of damp/moist paper towel in a petri dish lid. The lid was placed into an 8 oz Deli container (S-21216, Uline, Pleasant Prairie, WI, USA) with a lid until the eggs hatched (approximately 8-10 days). After hatching, crickets were transferred into cricket cages. Cricket cages consisted of 20 qt plastic storage containers (Gasket Box, Sterilite, Townsend, MA, USA) with four 100 mm circles cut on the side and two on the lid that were covered by screen mesh material.

### Cytometric genome size estimation of *A. domesticus*

The 1C (haploid) genome size of males and females of *A. domesticus* was estimated as described in [69]. In brief, a single *A. domesticus* head, a single *Drosophila virilis* female head (1C = 328 Mbp) and a portion of brain from a *Periplaneta americana* male (1C =3300) were placed together into 1 mL of ice-cold Galbraith buffer in a 2 mL Dounce tissue grinder. Nuclei were released by grinding with 10 strokes of an A (loose) pestle, then filtered through nylon mesh into a 1.5 mL microfuge tube. The DNA in the released nuclei were stained by adding 25 µl of propidium iodide (1 mg/mL) and allowed to stain 3 h in the dark at 4 ℃. The mean red PI fluorescence of the stained DNA in the nuclei of the sample and the standards was quantified using a CytoFlex flow cytometer (Beckman Coulter). Haploid (1C) DNA quantity was calculated as (2C sample mean fluorescence/2C standard mean fluorescence) times 328 Mbp for the *D. virilis* standard and times 3300 Mbp for the *P. Americana* standard. The estimates based on the two standards were averaged to produce a 1C estimate for each sample.

### Genome sequencing and assembly

A single adult male *A. domesticus* was shipped to Dovetail Genomics (now Cantata Bio, Scotts Valley, CA, USA). Genomic DNA was extracted and shipped to North Carolina State University and USDA ARS, Stoneville, MS for long read sequencing, and extracted nuclei were used for Chicago and Hi-C libraries.

Long read data was obtained using Pacific Biosciences technologies (PacBio, Menlo Park, CA, USA). Libraries were prepared using SMRTbell Express Template Prep Kit and Sequel Binding Kit 2.1 was used for polymerase binding. The binding reaction was loaded at 8 pmol concentration with a 15 h run time. The gDNA was sequenced on eight LR SMRTCells 1M v3 the Sequel I, for a total of 58.7 Gb of long read data. The genome was assembled from PacBio reads using CANU v1.8 [88]. The two sets of PacBio CLR reads were corrected separately and co-assembled. One set was iteratively corrected with the command canu -correct -p asm -d set1_round1 genomeSize=2.5g corMinCoverage=0 corMhapSensitivity=high corOutCoverage=100 -pacbio-raw set1/*.subreads.fastq.gz with the output of each round being the input for the next round. The reads were trimmed after three rounds of correction with the command canu trim -p asm -d set1_round3_trim genomeSize=2.5g correctedErrorRate=0.105 -pacbio-corrected set1_round3/asm.correctedReads.fasta.gz. The second set of reads was corrected with the command canu -correct -p asm -d set2_round1 genomeSize=2.5g corMinCoverage=0 corMhapSensitivity=normal corOutCoverage=100 -pacbio-raw set2/*subreads.fastq.gz and trimmed as set1. Lastly, the combined corrected reads were assembled with the command canu -assemble -p asm -d set1+set2 ‘genomeSize=2.5g’ ‘correctedErrorRate=0.105’ ‘batOptions=-dg 3 -db 3 -dr 1 -ca 500 -cp 50’ -pacbio-corrected set2_round1_trim/asm.trimmedReads.fasta.gz -pacbio-corrected set1_round3_trim/asm.trimmedReads.fasta.gz. This final assembly was polished with long read data in Arrow (SMRT Link 3.1.1, PacBio).

A Chicago library was prepared as described previously [89]. Briefly, ∼500ng of HMW gDNA was reconstituted into chromatin *in vitro* and fixed with formaldehyde. Fixed chromatin was digested with DpnII, the 5’ overhangs filled in with biotinylated nucleotides, and then free blunt ends were ligated. After ligation, crosslinks were reversed and the DNA purified from protein. Purified DNA was treated to remove biotin that was not internal to ligated fragments. The DNA was then sheared to ∼350 bp mean fragment size and sequencing libraries were generated using NEBNext Ultra enzymes and Illumina-compatible adapters. Biotin-containing fragments were isolated using streptavidin beads before PCR enrichment of each library. The libraries were sequenced on an Illumina HiSeq X to produce 208 million 2x150 bp paired end reads. A Dovetail HiC library was prepared in a similar manner as described previously [90]. Briefly, for each library, chromatin was fixed in place with formaldehyde in the nucleus and then extracted Fixed chromatin was digested with DpnII, the 5’ overhangs filled in with biotinylated nucleotides, and then free blunt ends were ligated. After ligation, crosslinks were reversed and the DNA purified from protein. Purified DNA was treated to remove biotin that was not internal to ligated fragments. The DNA was then sheared to ∼350 bp mean fragment size and sequencing libraries were generated using NEBNext Ultra enzymes and Illumina-compatible adapters. Biotin-containing fragments were isolated using streptavidin beads before PCR enrichment of each library. The libraries were sequenced on an Illumina HiSeq X to produce 143 million 2x150 bp paired end reads.

The Input *de novo* assembly, Chicago library reads, and Dovetail HiC library reads were used as input data for HiRise, a software pipeline designed specifically for using proximity ligation data to scaffold genome assemblies [89]. An iterative analysis was conducted. First, Chicago library sequences were aligned to the draft input assembly using a modified SNAP read mapper [91]. The separations of Chicago read pairs mapped within draft scaffolds were analyzed by HiRise to produce a likelihood model for genomic distance between read pairs, and the model was used to identify and break putative misjoins, to score prospective joins, and make joins above a threshold. After aligning and scaffolding Chicago data, Dovetail HiC library sequences were aligned and scaffolded following the same method.  Scaffolds that were determined to be “contaminants” (mitochondrial, duplicate, vector, or microbial as described in the next section) were removed prior to submission, resulting in 9,961 final scaffolds, and a total length of 2,346,604,983 bp. This Whole Genome Shotgun project has been deposited at DDBJ/ENA/GenBank under the accession JAHLJT000000000. The version described in this paper is version JAHLJT010000000.

To determine scaffolds that were from the metagenome, we used multiple tools to screen the scaffolds, including BBSketch and NCBI Refseq (https://www.biostars.org/p/234837/) and Kraken2 [92] in Omicsboxs (Biobam, Valencia, Spain, version 2.1.2, database RefSeq 2021.04, 2022.02). The scaffolds also were used in beta testing for GX, a screening tool for microbial contamination at NCBI (https://github.com/ncbi/fcs). We combined results from all three datasets (BBtools, Kraken2, GX) and manually annotated unclassified scaffolds with blastn [70], further using read depth to assign as insect or microbial. After this process, 80 scaffolds were removed as a result of NCBI screening tools, and 6,245 metagenomic scaffolds were submitted to NCBI MIMS as a metagenome/environmental, host-associated sample (accession number PRJNA908265).

### Annotation and analysis

Annotation of the *A. domesticus* genome assembly was by Dovetail Genomics. Repeat families found in the genome assemblies of *A. domesticus* were identified *de novo* and classified using RepeatModeler (version 2.0.1) [93]. RepeatModeler depends on RECON (version 1.08) and RepeatScout (version 1.0.6) for the *de novo* identification of repeats within the genome. The custom repeat library obtained from RepeatModeler was used to discover, identify, and mask the repeats in the assembly file using RepeatMasker (Version 4.1.0) [94]. Coding sequences from *Locusta migratoria*, *Teleogryllus occipitalis*, *Laupala kohalensis*, and *Xenocatantops brachycerus* were used to train the initial *ab initio* model for *A. domesticus* using AUGUSTUS (version 2.5.5) [95], with six rounds of prediction optimization. The same coding sequences also were used to train a separate *ab initio* model for *A. domesticus* using SNAP (version 2006-07-28) [89]. RNAseq reads were mapped to the genome using STAR aligner (version 2.7) [96], and intron hints were generated with the bam2hints tools within AUGUSTUS. MAKER, and SNAP (intron-exon boundary hints provided from RNA-Seq) and were then used to predict genes in the repeat-masked reference genome. Swiss-Prot peptide sequences from the UniProt database were downloaded and used in conjunction with the protein sequences from the same training species to generate peptide evidence in Maker [97]. Only genes that were predicted by both SNAP and AUGUSTUS were retained in the final gene sets. AED scores were generated for each of the predicted genes as part of the MAKER pipeline to assess the quality of the gene prediction. Genes were further characterized for putative function by performing a BLAST [70] search of the peptide sequences against the UniProt database. tRNA were predicted using the software tRNAscan-SE (version 2.05) [98]. Predicted genes were analyzed by BUSCO (v.2.0) [99], using the lineage dataset insecta_odb9 (Creation date: 2016-10-21, number of species: 42, number of BUSCOs: 1658). Predicted genes and sequence information are in supplemental files (Files S10-11).

Genes annotated for CRISPR-Cas9 (*A. domesticus vermilion, Adv*, and *actin*, *Ad_muscle_actin*) were submitted to NCBI as accession #2652422.

We annotated repetitive elements against the publicly available genomes consisting of six crickets and one locust. The sources of the genomes used in the analysis are as follows: *Apteronemobius asahinai*, GCA_01974035.1; *Gryllus bimaculatus*, GCA_017312745.1; *Laupala kohalensis*, GCA_002313205.1; *Teleogryllus oceanicus*, www.chirpbase.org, *Teleogryllus occipitalis*, GCA_011170035.1; *Locusta migratoria*, GCA_000516895.1. Libraries of repetitive elements were compiled by RepeatModeler-2.0.3 [93] for each species. Resulting libraries were classified using reference-based similarity search in RepBase (20170127 edition). RepeatMasker-4.1.2-p1 [94] was used with alignment option ‘-a’ to annotate the repetitive elements in the genome assemblies. The resulting alignment files were processed by the scripts with RepeatMasker pipeline and our in-house scripts (https://github.com/kataksk/repeatlandscape_parser.git). To analyze correlation between the repetitive elements and intron lengths, R packages GenomicFeatures [100] was used to obtain gene structures of gene models.

To extract virus sequences from the genome, blastn was performed against *Chilo iridescent* virus (GenBank: AF303741.1). Quality assessment of the extracted virus-like scaffolds was performed by CheckV (v1.0.1) [101].

### CRISPR target gene for knock-out/in

We chose the 5’ of the *Ad vermilion* gene as our CRISPR target site for knock-in experiments. We identified the *Ad vermilion* gene from the *A. domesticus* genome assembly using a transcriptome mRNA sequence (not complete, missing 3’) by Blastation (TM Software, Inc., Arcadia, CA, USA). We were able to identify eight exons including the first exon (Figure 5a). We then used the second exon (around 150 bp) to select three target CRISPR targeting sites for designing sgRNAs (Table S9) using GGP sgRNA designer (https://portals.broadinstitute.org/gpp/public/analysis-tools/sgrna-design). The target sgRNA sequences were ordered from Synthego (Redwood City, CA, USA). To verify the knock-out sequence in *A. domesticus*, genomic DNA extraction was using a ZymoResearch Quick-DNA insect miniprep kit (Irvine, CA, USA), forward primer “GAGCAGTAGGCGAGAAAG” and reverse primer “TCCAAACGCAGAAGAGACCA” to amplify gDNA. After PCR, we used primer “TGTAGTGAGTGTTATCGCCA” to sequence the PCR product using the Oklahoma Medical Research Foundation sequencing service facility (OMRF DNA sequencing, https://omrf.org/). Using these data we compared the gRNA sequence to wild type sequence to determine the indel(s) generated by our knock-out and knock-in experiments (Table 3).

### Knock-in DNA construct design

Fo knock-in construct design we used the *Ad muscle actin* gene promoter to express the marker gene, enhanced green florescent protein (EGPF) via CRISPR/Cas9. We utilized the muscle actin gene from *Tribolium castaneum* and *Drosphila melanogaster* to conduct a BLAST search of the *A. domesticus* transcriptome data and used the mRNA sequence from that search to identify the *Ad muscle actin* gene in the genome assembly. We used 1.5 kb upstream from the start codon of the *Ad muscle actin* genomic sequence and searched for a promoter region using the Neural Network Promoter Prediction tool (https://fruitfly.org/seq_tools/promoter.html).

To design the DNA construct, 455 bp of the *Ad muscle actin* promoter sequence with its 5’UTR were placed upstream of the EGFP coding sequence along with a sv40 polyadenylation signal 3’ of EGFP as the marker gene. A section of 86 bp of gDNA included CRISPR knock-in site(s) placed on both sides of the marker gene as our final knock-in DNA construct, Ad-actin-EGFP-KI. This DNA sequence was synthesized by GeneScript (Piscataway NJ, USA) in their pUC57 vector plasmid as the final product.

### Microinjection solution preparation

The concentrations of the components of knock-out or knock-in microinjection solutions were: 100 ng/μL of DNA construct Ad-actin-EGFP-KI (only for knock-in treatment), 10 pmol/ μL of sgRNA, 1 µg/μL of TrueCut Cas9 protein V2 (ThermoFisher, Waltham, MA, USA), and 20% phenol red buffer (Sigma Aldrich, St. Louis, MO, USA). The solution was mixed and incubated at room temperature for 5-10 min for Cas9-sgRNA binding, and the solution was kept on ice during microinjection. For knock-out experiments we used either three sgRNAs or sgRNA#1 alone for microinjections (Table S9, sgRNA target sequences). For knock-in experiments, we only used sgRNA#1 in the microinjection solution.

### Cricket egg microinjection protocol for CRISPR

An egg lay dish with hydrated water crystals was placed into an adult wild type cricket colony for 4 h to receive fresh and non-developed eggs. Eggs were collected and washed using the protocol described above in the “House Cricket Colony Maintenance” section. After washing the eggs from the water crystals, a solution of 70% ethanol (ethanol/deionized water) was used to briefly rinse the eggs for 10-15 s and quickly rinsed with deionized water at least 3 times. The eggs were placed on microinjection slides (Figure S12). To prepare the microinjection slide , a piece of black filter paper was cut into 50 x 10 mm pieces and laid on a standard clean glass microscope slide (75 x 26 mm). Another two pieces of black filter paper strips, 50 x 2 mm, were cut and laid one on the top of the other placed in the middle of the larger piece of black filter paper. Deionized water was added to the filter paper, and eggs were placed on the edge of two sides of the filter paper strips for microinjection. The wet filter prevented the eggs from drying out during microinjection and allowed a backstop to keep the eggs from moving during the procedure. The black paper made visualizing the eggs easier compared to using white filter paper. The needle for microinjection was pulled fro, Standard Glass Capillaries (1B100-4, World Precision Instruments) using a P-2000 needle puller (Sutter instruments). The following puller apparatus settings were used: Heat: 335 Fil: 4 Vel: 40 Del: 240 Pul: 120. For microinjection, 1 μl of microinjection solution was loaded into the back of the needle with Femtotips (Eppendorf, Hamburg, Germany) and all air bubbles were removed from the liquid in the needle, particularly the tip. The filled needle was then applied to a FemtoJet 5247 microinjector (Eppendorf) and air pressure of 4-7 psi was used for holding the solution in place and 9-10 psi for injection. Before injection, forceps were used to gently break the needle tips to achieve a small opening. The injection site for eggs was 2/3 from the smaller end. After injection, the black filer paper containing the injected eggs was transferred from the glass slides to a piece of wet paper towel and kept in a 150 x 10 mm petri dish with lid. The dish was placed into a hatch chamber (Modular Incubator Chamber, MIC-101, Billups-Rothenberg, San Diego, CA, USA) with two wet paper towels for 10-12 d.

### Screening for CRISPR knock-out/in and established colonies

To screen for either white eye (*Ad vermilion* knock-out) or EGFP (EGFP knock-in) expression phenotypes, freshly hatched G0 crickets were transferred from their egg incubation chamber (described above) into a standard petri dish with lid. The petri dish was then placed on top of ice to immobilize them by cooling them down for three minutes. The crickets where then observed under a fluorescent dissecting microscope with digital camera attached (Leica M125, Leica Microsystems Inc., Deerfield, IL, USA) for either different eye coloration from wild type in knock-out treatments (using a standard white LED ring light) or using a blue florescent light and GFP filter to check for knock-in EGFP expression phenotype. Photos were taking using a QImaging Retiga R6 digital camera attached to the microscope. After screening, G_0_ crickets with positive knock-out white/vermilion eye (full or partial) phenotype were separated and reared to adulthood in containers described in the house cricket colony maintenance section above. Crickets with wild type eye phenotype from knock-out experiments were not kept. G_0_ crickets with EGFP positive phenotype were reared in separate containers from those negative for EGFP until adulthood from the knock-in experiments. Once these crickets became adults, self-cross experiments were set up (two males with four to six females) groups for positive G_0_s from the knock-out experiments. For knock-in experiments, small out-cross groups were set up using EGFP positive G_0_s with two G_0_s out-cross to 2-3 wildtype crickets, and one self-cross group from all EGFP negative G_0_s pooled together in the same cage. Cross groups were maintained and eggs collected as described in the house cricket colony maintenance section above. Once the G1 generations began to hatch from eggs, we then screened and separated all white eye color phenotype knock-out G_1’s_ from self-cross groups, and EGFP positive G_1_s from out-cross groups and established separate colonies from all crosses. No EGFP positive G1 crickets were found from the pooled EGFP negative G0 injected cricket colony.

### RNAi target genes and dsRNA sequences

Short portions of the *Ad vermilion* mRNA gene sequence (679 bp) were used to design 450 bp (51-500 bp) dsRNA (dsAdV) (Table S9) to target the *Ad vermilion* gene for knock-down via RNAi. An additional dsRNA (dsEGFP) also was made using 450 bp of EGFP coding sequence (region 96-545 bp) and was used both to knock-down EGFP in our knock-in cricket lines via RNAi as well as a negative control for the RNAi experiments for knocking down *Ad vermilion* for white eye color via RNAi. Both dsRNA sequences were ordered from Genolution Inc. (Seoul, Korea).

### Cricket microinjection and screening for RNAi

The concentrations of components of the cricket RNAi microinjection solution were: 2.5 µg/ul for dsRNA (dsAdV or dsEGFP) with 20% phenol red buffer. Two separate experiments were utilized to demonstration of RNAi efficacy in *A. domesticus*; one using wild type crickets and the other using our genetically modified strain. The dsAdV was injected into wildtype crickets to knock down eye color. The dsEGFP was injected into wildtype crickets both as a negative control and also in separate experiments to knock down EGFP expression in our AdVELV1-3 EGFP expression strain. Two to three sets of microinjections of each RNAi experiment were conducted using young nymphs (following their first molt after hatching from eggs). Crickets were placed on ice for 5 min to immobilize them and were subsequently injected in the posterior abdomen (Figure S12). Around 25 to 30 crickets were injected using a total of 4 µl of injection solution (approximately 0.15 µL per injection). Following injection, crickets were maintained in a medium sized cage (Kritter Keeper®, 4.80 x 7.40 x 5.60 in, Lee’s Aquarium & Pet Products, San Marcos, CA, USA) with food and water. After microinjection, crickets were screened based on eye color or EGFP expression every week and phenotype changes were documented.

## Acknowledgments

We thank Ken Friesen for bioinformatics support, and Eric Tvedte for analysis of *A. domesticus* scaffolds in GX beta testing. Dr. Dossey also thanks Mr. Tyler Tatum (Ripple Technology, LLC, Atlanta, GA, USA) for his help and organizational support. This material is based upon work supported by the Defense Advanced Research Projects Agency (DARPA) under Contract No. 140D6318C0055. The research was completed under Cooperative Research and Development Agreement Number 58–3020–7–013 between ARS, ATB, and NCSU. This study was supported by the Cabinet Office, Government of Japan, Cross-ministerial Moonshot Agriculture, Forestry and Fisheries Research and Development Program, “Technologies for Smart Bio-industry and Agriculture” (BRAIN) [JPJ009237 to KK]. This research was performed in accordance with Kansas State University Research Compliance Office, Institutional Biosafety Committee registration number 1191, “Functional Genomics of Stored Product Insects”. This work was supported, in part, by the Intramural Research Program of the National Human Genome Research Institute, National Institutes of Health (SK). This genome assembly is a contribution to the i5k and USDA ARS AgPest100 projects. This work utilized the computational resources of the NIH HPC Biowulf cluster (https://hpc.nih.gov). Mention of trade names or commercial products in this publication is solely for the purpose of providing specific information and does not imply recommendation or endorsement by the U.S. Department of Agriculture. USDA is an equal opportunity provider and employer.

## Supporting information

**S1 Table. Genome size of *A. domesticus***. 1C = the amount of DNA in a gamete (1C is an average of the gametes with and without the X in the male).

S2 Fig. Comparison of the lengths of BUSCO reference genes (bp) in *Tribolium castaneum* and *Acheta domesticus*.

**S3 File. Metagenomic scaffolds from the *A. domesticus* genome assembly.** Spreadsheet “Ado_metagenome_Kraken2”) details the output from files tentatively identified as non-insect through the first screening; “Blastn_unclassified” is the output from blastn of the unclassified scaffolds to NCBInr; “Metagenome_summary” are the scaffolds submitted to NCBI as *A. domesticus*-associated sequences.

**S4 File. Screen of viral genome sequences through CheckV (Nayfach, et al., 2021).** Data is sorted according to high, low, or medium quality, or not-determined based on matches to the CheckV database.

**S5 File. Immune-related transcripts that are predicted from the annotation of the *A. domesticus* assembly.** Predicted transcripts include: PGRP – peptidoglycan-recognition protein; GNBP - β-1-3-glucan-binding protein; lysozyme; PPO – prophenyloxidase.

S6 Table. Comparison of intron length, genome size, and repeat content in orthopteran species (six crickets, including *Acheta domesticus*, *Apteronemobius asahinai*, *Gryllus bimaculatus*, *Laupala kohalensis*, *Teleogryllus occipitalis*, *Teleogryllus oceanicus* and one locust, *Locusta migratoria*). A). Correlation between total repeat content and genome size; B) Correlation between total repeat content and median length of intron; C) Correlation between genome size and median length of intron; and Comparison of D) LTR, E) LINE, and F) DNA transposable elements.

S7 Fig. Supporting data for the comparison of intron length, genome size, and repeat content in orthopteran species (six crickets, including *Acheta domesticus*, *Apteronemobius asahinai*, *Gryllus bimaculatus*, *Laupala kohalensis*, *Teleogryllus occipitalis*, *Teleogryllus oceanicus* and one locust, *Locusta migratoria*). A) LINE, B) DNA, and C) LTR transposon subclasses and correlation with intron length and genome size.

S8 Table. CRISPR sgRNA target sequences for the *Ad vermillion* gene.

S9 Table. Double-stranded RNA sequences used for RNAi experiments.

S10 File. Name and annotation of predicted *A. domesticus* genes, with scaffold number and length.

S11 File. Transcript sequences of predicted *A. domesticus* genes.

**S12 Fig. *A. domesticus* egg microinjection slide.** A) Slide set up with eggs positioned on each side of the filter paper strips; B) Injected eggs showing microinjection site with phenol red color in eggs; and C) Developed eggs after microinjection showing eye pigmentation.

